# Intra-tumor heterogeneity, turnover rate and karyotype space shape susceptibility to missegregation-induced extinction

**DOI:** 10.1101/2021.11.03.466486

**Authors:** Gregory J. Kimmel, Richard J. Beck, Xiaoqing Yu, Thomas Veith, Samuel Bakhoum, Philipp M. Altrock, Noemi Andor

**Author notes:** Authors contributed equally to this work (Gregory J. Kimmel, Richard J. Beck).

## Abstract

The phenotypic efficacy of somatic copy number alterations (SCNAs) stems from their incidence per base pair of the genome, which is orders of magnitudes greater than that of point mutations. One mitotic event stands out in its potential to significantly change a cell’s SCNA burden–a chromosome missegregation. We present a general deterministic framework for modeling chromosome missegregations and use it to evaluate the possibility of missegregation-induced population extinction (MIE). The model predicts critical curves that separate viable from non-viable populations as a function of their turnover- and missegregation rates. Missegregation- and turnover rates estimated from a PAN-cancer scRNA-seq dataset of 15,464 cells are then compared to predictions. The majority of tumors across all cancer types had missegregation- and turnover rates that were within viable regions of the parameter space. When a dependency of missegregation rate on karyotype was introduced, karyotypes associated with low missegregation rates acted as a stabilizing refuge, rendering MIE impossible unless turnover rates are exceedingly high. Intra-tumor heterogeneity, including heterogeneity in missegregation rates, increases as tumors progress, rendering MIE unlikely.

**Author Summary:** When a cell missegregates a chromosome while dividing, the chance is high that its two daughter cells will behave drastically different from each other and from their parental cell. Chromosome missegregations are therefore one of the most powerful forces of phenotypic diversity. We developed a mathematical model of chromosome missegregations that allows for this cell-to-cell diversity to be accounted for. The model serves to help understand how selection acts upon cells with versatile chromosome contents, as a tool for genotype-to-phenotype mapping in various microenvironments. As a first application example we used the model to address whether there exists an upper limit on missegregation rate, beyond which cancer populations collapse. Chromosome missegregations are common. They occur in 1.2-2.3% per mitosis in normal cells [1] and in cancer cells their rate is between one and two orders of magnitudes higher [2]. The model revealed that the upper limit of missegregation rate is a function of the tumor’s turnover rate (i.e. how fast the tumor renews itself). In heterogenous populations however, cells with low missegregation rates protect the population from collapse. Intra-tumor heterogeneity, including heterogeneity in missegregation rates, increases as tumors progress, rendering missegregation-induced extinction unlikely.

## 1 Introduction

Aneuploidy, defined as a chromosome number that is not the exact multiple of the haploid karyotype, is common across several cancers, including non-small-cell lung, breast, colorectal, prostate cancer and glioblastoma [3–7]. The main driver of aneuploidy is chromosomal instability (CIN). CIN-induced genomic changes can be subdivided into two categories: the whole gain or loss of a chromosome (numerical CIN) or changes within localized regions of a chromosome (structural CIN).

Thompson and Compton used live cell imaging to evaluate the fidelity of chromosome segregation, finding missegregation rates ranging from 0.025-1 % per chromosome per mitosis [2]. We distinguish between unpredictable and predictable factors governing a cell’s risk to missegregate. Unpredictable events include DNA double-strand breaks (DSBs). Their location in the DNA appears to be random, yet has been shown to influence the likelihood of mitotic delay and subsequent missegregation events [8–11]. This delay allows for DNA damage response (DDR) during mitosis and thus protects the genome from structural damage, but at the expense of increasing risk for numerical instability [12]. Predictable factors that increase the incidence of missegregations include high ploidy [13] and suboptimal kinetochore-microtubule attachment stability. Tetraploid cells are more likely to fail to cluster centrosomes into two poles, leading to multipolar division. While multipolar divisions are likely lethal, multipolar mitosis can also cause the poles to coalesce leading to a pseudobipolar division and chromosome missegregations ([14]). Kinetochore-microtubule attachment stability must fall within a narrow permissible window to allow for faithful chromosome segregation [12, 15–17] and is influenced by both *cell-intrinsic* [12] and *extrinsic* factors. An example of extrinsic factors are Vinca alkaloids (e.g. vincristine, vinblastine), – a class of cytotoxic drugs which act directly upon the microtubule network [18], causing increased missegregation rates [19]. But even cytotoxic drugs not directly targeting the microtubule network have been shown to significantly impede segregation fidelity, through aforementioned stimulation of DDR and mitotic delay [20]. Drugs targeting the DDR are likely to induce numerical instability [20], suggesting that DNA-damaging therapies impart part of their cytotoxicity by interfering with chromosome segregation fidelity [21]. Changes of the tumor microenvironment such as glucose deprivation, hypoxia and acidification have also been linked to CIN [22, 23].

Anueploidy and CIN are often coupled and can create a positive-feedback loop in which further structural or whole chromosome aberrations accumulate over time [24, 25]. But normal cells do not tolerate missegregations – aneuploid daughter cells are immediately cleared from the cell pool through apoptosis during G1 following a missegregation [26, 27]. The sudden genome-dosage imbalance caused by a missegregation event induces p53-mediated cellular senescence in the subsequent G1 phase [13, 26–28]. While cancer cells evolve to more proficiently avert missegregation-induced cell death, missegregations still activate p53 in the G1 phase, even among cancer cells [28], albeit less reliably [29]. High levels of CIN have been observed to be tumor suppressive in breast [30], ovarian, gastric, and non–small cell lung cancer [31]. The above suggests that a non-monotonic relationship between cell fitness and CIN likely exists, with a threshold of a critical level of CIN (which may be cancer type specific). A possible therapeutic avenue to target and exploit the degree of CIN in patients is therefore guided by the premise that a Goldilocks window exists for cancer to thrive.

Gusev et al. [32] modeled the evolution of cell karyotypes via chromosome mis-segregations as a random branching walk. Using this model, the authors estimated the fraction of clones surviving as a function of mis-segregation rate and approximated a theoretical limit for mis-segregation rate for a diploid population to survive without a complete loss of any chromosome type. In a follow-up publication, the same authors used a semianalytical approach to analyze the asymptotic behavior of this model, simulating evolution of the copy number of just a single chromosome type [33]. They compared various mechanisms of chromosome missegregations with respect to their ability to generate a stable distribution of chromosome numbers. Elizalde et. al. explored the phenotypic impact of CIN using a Markov-chain model and confirmed the existence of optimal chromosome missegregation rates [34]. The authors assumed that cells were not viable if they contained nullisomy (the loss of all copies of a chromosome), paired with a corresponding upper limit of eight copies. These assumptions were justified through a sensitivity analysis [35]. The main conclusion of the paper established that missegregation rates drove heterogeneity more than the age of the tumor. Under what circumstances missegregations lead to tumor extinction however remains unclear. Here we derive necessary conditions that drive a tumor population to nonviable karyotypes, typically through either nullisomy or an upper limit on the number of sustainable copies per chromosome (e.g. eight [34]). We further refer to these conditions as missegregation-induced extinction (MIE).

The remainder of this manuscript is structured as follows. We first motivate the existence of MIE with a phenomenological equation of ploidy movement that takes the form of a diffusion-reaction equation. A heuristic argument comparing the time scales of net growth and missegregation will imply the existence of turnover and missegregation rates that allow for MIE. We then develop our mathematical framework–a general coupled compartment model of chromosome mis-segregations. This model is simplified to a version more amendable to theoretical analysis, while still retaining the qualitative behavior and form. Theoretical results are derived and presented on the existence of MIE and when it can be evaded. Next, we derive turnover and missegregation rates from a PAN-cancer scRNA-seq dataset from 14 tumors and quantify their relationship to ploidy and to the copy number of individual chromosomes. Finally, we use the scRNA-seq derived measurements to predict which tumors are most sensitive to MIE.

### Key terms and definitions

- A chromosome **mis-segregation** segregates chromosomes asymmetrically among daughter cells, causing an aneuploid state.
- **Aneuploidy**:= a chromosome number that is not a multiple of the haploid complement. An **euploid** cell carries a multiple of the same set of chromosomes. Haploids (1 copy of each chromosome), diploids (2 copies) and polyploids (> 2 copies) are all examples of euploidy.
- **Karyotype**:= a numeric vector with one entry per chromosome **type** (e.g. we have 22 types when considering the set of human autosomes). Each entry indicates the **copy number** of that type of chromosome. Each karyotype corresponds to a compartment of a multi-compartment ODE.
- **Ploidy**:= the total number of chromosomes occurring in the nucleus of a cell. Also referred to as **DNA content** in the continuous setting.
- **Karyotype composition**:= the relative frequency of all karyotypes in a population at a given time.
- **Set of viable karyotypes**:= all possible karyotypes associated with non-negative growth rates. This can either be a single **interval** (e.g. 22 - 88 chromosomes) or multiple intervals of karyotype viability interspersed between nonviable regions.

## 2 Results

### 2.1 Cell turnover rates and susceptibility to MIE

Consider a simple birth-death process on ploidy space, where, for the moment, we are interested only in the total amount of DNA content of a cell. If only missegregations facilitate the movement in DNA content during mitosis, one can crudely approximate the total population *n*(*p*) as a function of DNA content *p* by the following partial differential equation (PDE):

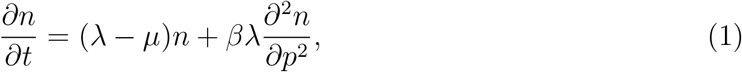

where λ, *μ* are the birth, death rates, respectively and *β* is the missegregation rate. For boundary conditions, we make the assumption that there exists *p* ∈ (*p*_min_, *p*_max_), such that for DNA content outside this range, the population cannot survive. We also assume that *r* = λ – *μ* > 0, that is, in the absence of missegregation, this population is favored to grow.

We now appeal to a heuristic argument of time scales. Let *T_p_* be the timescale on which missegregation events happen, which, in Fickian diffusion is proportional to: 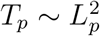, where *L_p_* is the characteristic amount of DNA content shifted during a missegregation event. Because only dividing cells mis-segregate, we have: 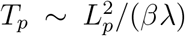. Similarly, let *T_r_* be the time scale on which the cell population grows: *T_r_* = 1/*r*. If *T_p_* ≪ *T_r_* then extinction via missegregation (hereby “missegregation-induced extinction” or MIE) is possible. The inequality implies that MIE can occur if

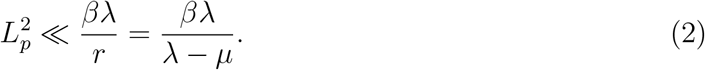

An important takeaway from this simple argument is that the characteristic scale *L_p_* can play a significant role and is tied to the typical change in DNA content when a missegregation occurs. The narrower the interval of viable DNA content, the weaker the condition, i.e. more combinations of turnover and missegregation rate will exist that lead to MIE.. Second, it is interesting to see that in theory, one does not need to increase missegregation, but rather can also increase birth and death rates in such a way that the quantity (1 — *μ*/λ) decreases. This suggests that tumors with high turnover rates may be more susceptible to MIE.

### 2.2 General discrete model of chromosome mis-segregations

Equation (2) was derived under the assumption that shifts in DNA content happen on a continuous scale. But in reality chromosomes are discrete units of information. To investigate whether the impact of turnover rate on MIE remains valid in the discrete setting, we developed a general compartment model that describes the evolution of populations by their karyotypes. Let the *M*-dimensional vector 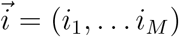 contain the number of copies *i_k_* ≥ 0 of the *k*th component. An example would be looking at the copy number of whole chromosomes (i.e. *M* = 23). Alternatively one can parse the chromosome into the *p*, *q* arm (i.e. *M* = 46). Movement between the compartments occurs via missegregation. We encapsulate this information in the tensor **q** with non-negative components 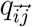, which is the probability with which a cell from compartment 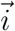 moves to compartment 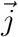 (Fig. 1). We require a conservation of copy number (which will hold for any resolution). Let 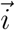 be the parent and 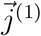 and 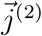 be the offspring, then it must be that

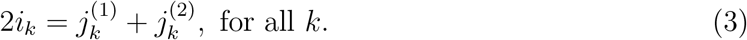

**Figure 1:**
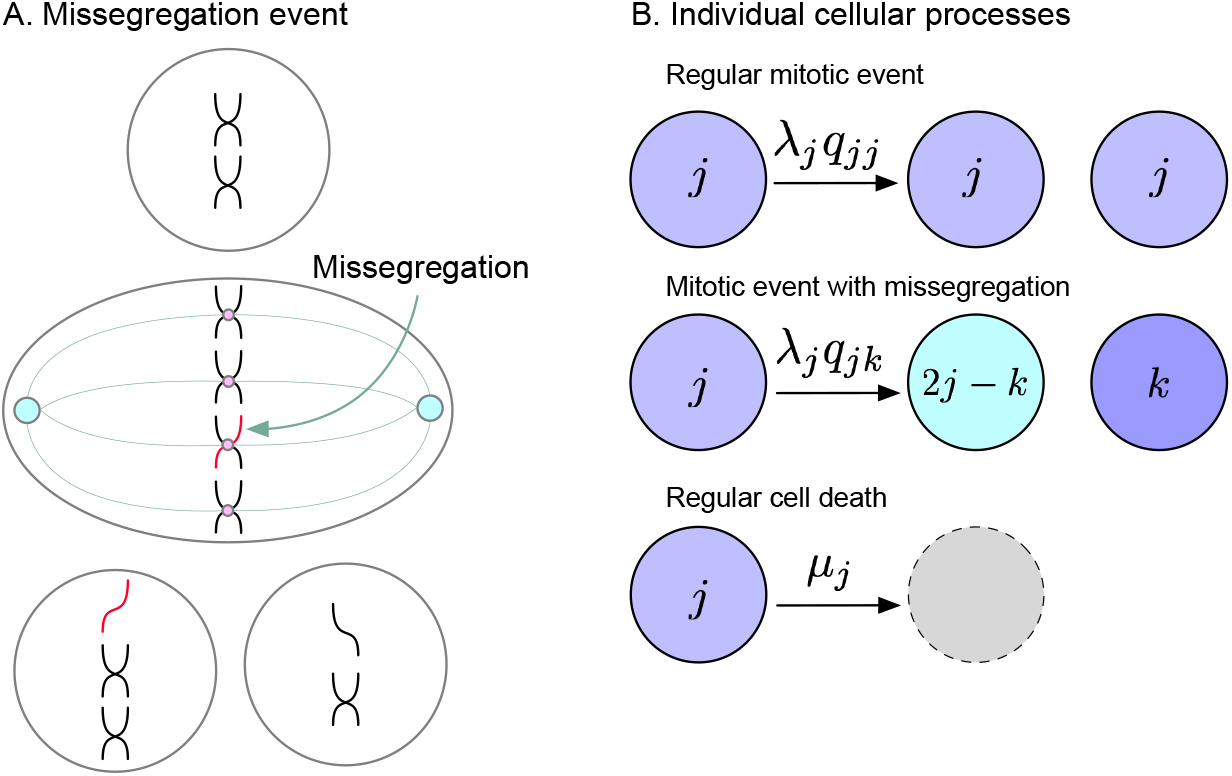
Mathematical modeling of chromosome missegregations. (**A**) Missegregation event. During anaphase, one daughter cell improperly takes both chromosomes leading to aneuploidy. Note the copy number conservation assumption here. (**B**) Model’s individual cellular processes. The tensor *q_jk_* encodes the probability at which a cell with copy number *j* may produce offspring with copy number *k* (and also 2*j* – *k* by copy number conservation). Thus karyotype *j* goes through anaphase with faithful chromosome segregations at a rate λ*_j_q_jj_* and missegregates into karyotype *k* (and 2*j* – *k*) at a rate λ*_j_q_jk_*. Cell death occurs with rate *μ_j_*.

This imposes structure on **q** since 0 ≤ *j_k_* ≤ 2*i_k_*, we must have 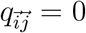 if *j_k_* > 2*i_k_* for any *k*, thus **q** will be sparse for many applications.

We are now in a position to write a general *M*-dimensional birth-death process:

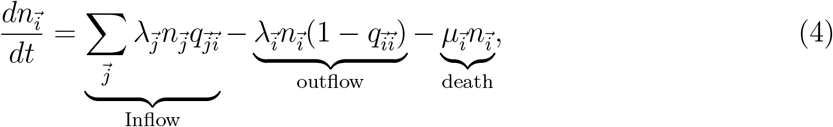

where 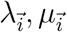 are the state-dependent birth and death rates, respectively. Equation (3) enters into (4) with the flow rate λ**q**. Model parameters are summaryzied in Table 1.

**Table 1:**
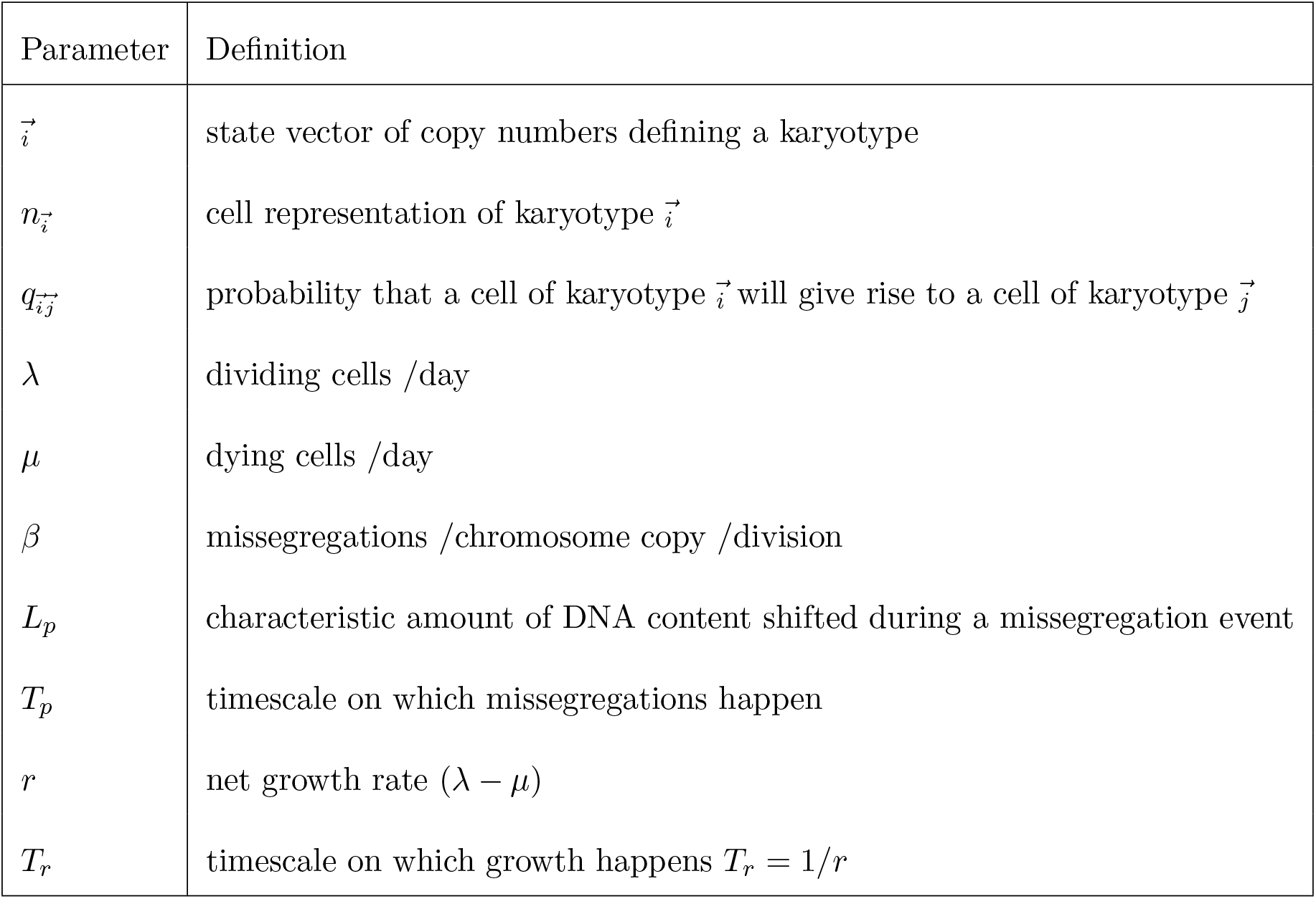
Common model parameters used throughout the manuscript.

| Parameter | Definition |
| --- | --- |
| $\vec{i}$ | state vector of copy numbers defining a karyotype |
| $n_{\vec{i}}$ | cell representation of karyotype $\vec{i}$ |
| $q_{\vec{i}\vec{j}}$ | probability that a cell of karyotype $\vec{i}$ will give rise to a cell of karyotype $\vec{j}$ |
| $\lambda$ | dividing cells /day |
| $\mu$ | dying cells /day |
| $\beta$ | missegregations /chromosome copy /division |
| $L_p$ | characteristic amount of DNA content shifted during a missegregation event |
| $T_p$ | timescale on which missegregations happen |
| $r$ | net growth rate ($\lambda - \mu$) |
| $T_r$ | timescale on which growth happens $T_r = 1/r$ |

We note that in the absence of missegregation, we require 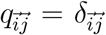 where we are using the vector Kronecker delta that is 1 if 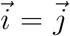 and 0 otherwise. This uncouples equation (4) to the classic deterministic birth-death process *dn_i_/dt* = (λ_*i*_ – *μ_i_*)*n_i_* as expected.

We define the shift, *t*, as the net difference in the copy number of a given chromosome type *k*, between the parental cell and its daughter cells. *t* can be positive or negative and accounts for the fact that missegregations can partly or entirely compensate each other. The probability *P*(*t|i_k_*), that a cell with *i_k_* chromosome copies will have a daughter cell with a shift of *t* copies has been derived by Gusev et al. [33] as:

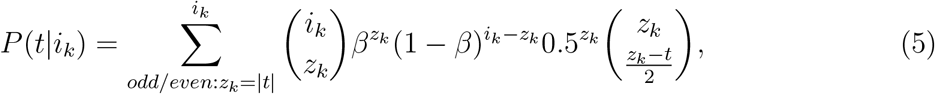

where missegregations are assumed independent and *β* is the missegregation rate per chromosome copy per division. If *t* = 0 this means that both daughter cells have the same copy number for chromosome *k*. If *t* ≠ 0, then there is one cell with *i_k_* + *t* copies and one cell with *i_k_* – *t* copies. We can thus calculate the probability of transitioning from karyotype 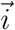 to 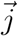 as:

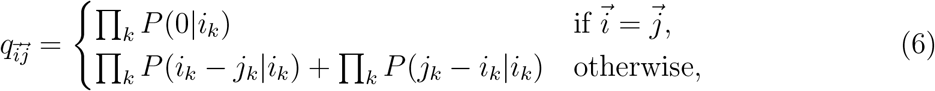

Note there is no inconsistency, if 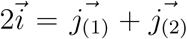, then 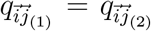, which satisfies the copy number conservation. Further, if we let β → 0 we would have:

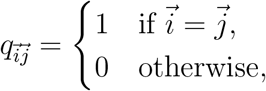

which we recognize as the Kronecker delta defined above.

In contrast to the continuum model given in equation 1, this model takes into account that chromosomes are discrete units of information. We implemented this model in R, allowing numerical simulations of karyotype evolution under variable initial conditions and biological assumptions. Finer genomic resolution leads to more compartments, thereby increasing computational resources required for numerical solutions. The remainder of this manuscript will use the resolution of whole chromosomes to define a karyotype. When missegregation, death and birth rates are independent of karyotype, we will refer to them as *homogeneous*. Conversely, *intra-tumor heterogeneity* in either of these rates will be modeled as a dependency on karyotype. Homogeneous rates imply that these rates will always stay constant over time. Constant rates however do not imply homogeneity within the population, since a heterogeneous but stable karyotype composition will appear constant despite representing multiple rates. In summary, this is a flexible framework, offering the possibility to model a variety of biologically relevant dependencies (e.g. missegregation rate can vary across karyotypes) and variable genomic resolutions.

### 2.3 Chromosomal aggregate model

The model given by equation (4) is complicated and cumbersome. A simpler model, amendable to analysis involves aggregating all chromosomal data into one index. Alternatively, it can be thought of as focusing on a dosage-sensitive chromosome, that must be present at copy numbers between one and five in order for a cell to survive. We note that this implies that the existence of all but one chromosome is negligible; hence all three terms, “karyotype”, “copy number” and “ploidy” become equivalent. Mathematically, there are many ways to collapse our *M*-dimensional model to 1D, and one such way is to just sum over all indices to get the aggregated number of copies:

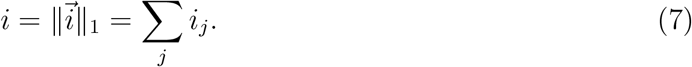

Then our system is given by:

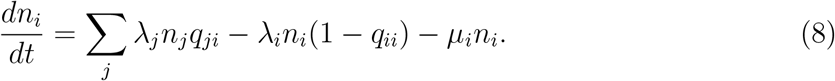

We will further suppose that the parameters of the model are not dependent on karyotype prevalence (e.g. λ_*i*_ is not dependent on any *n_i_*, such as through a carrying capacity or Allee effect etc). This allows us to easily write the Jacobian, which is simply the coefficients of *n_j_* in equation (8)

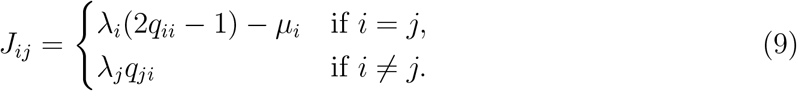

Let [*k*, *K*] with *k*, 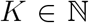 be the interval (not necessarily finite) of viable karyotypes of the aggregate model (eq. (8)). The Jacobian (9) contains information on the local behavior of the system near the extinction state *n_i_* = 0 for all *i*. If all the eigenvalues of the Jacobian at the extinction point are negative, then MIE occurs. The critical curve that separates MIE from exponential growth is when the maximum eigenvalue of *J* is 0.

### 2.4 Ruling out MIE

Here, we establish sufficient conditions for MIE to not occur based on Gershgorin’s circle (GC) theorem [36]. The theorem bounds the locations of the eigenvalues in the complex plane for a given matrix A, with elements *a_ij_*. The GC theorem stipulates that the eigenvalues must be contained in the circles with centers *a_ii_* and radii *R* = ∑_*i*≠*j*_|*a_ij_*|.

Since MIE can be evaded if the maximum eigenvalue exceeds 0, a sufficient condition is that none of the GCs contain a part of the negative reals. Table 2 describes sufficient conditions to avoid MIE for various biological assumptions.

**Table 2:**
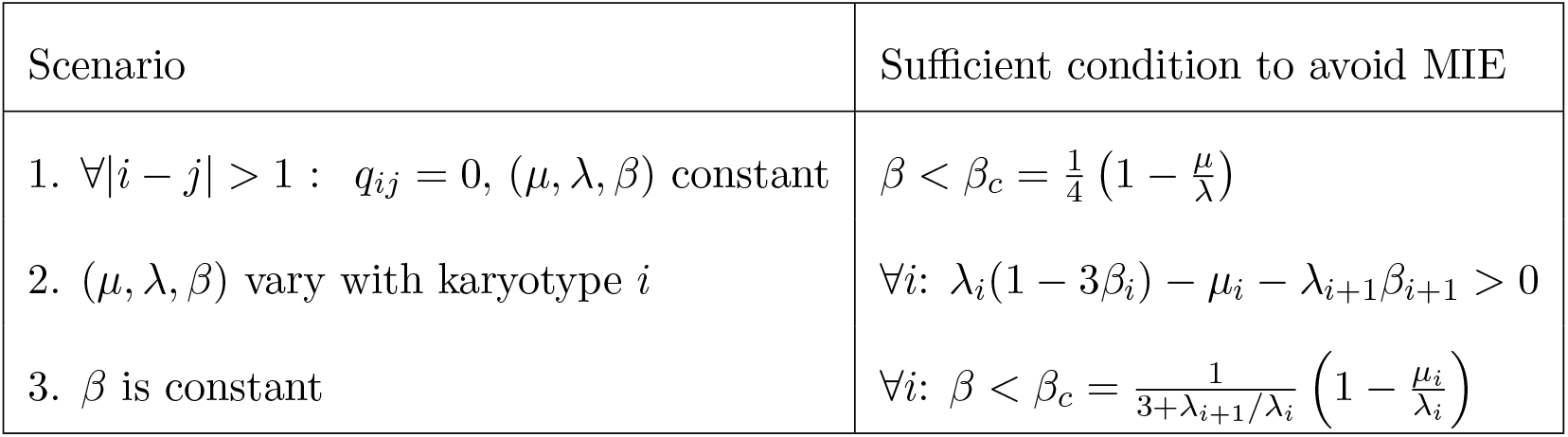
Sufficient conditions to avoid MIE for various biological scenarios: 1) Only ≤ 1 chromosome can missegregate per division; all rates are independent of karyotype (i.e. homogeneous). 2) Heterogeneous birth-, death-, and/or missegregation rates. 3) Homogeneous missegregation rate.

The general problem for arbitrary **q** can be handled numerically, but analytical conclusions can only be made for specific forms of **q**. The conditions required are given by finding when the GC’s are all contained in the positive half-plane:

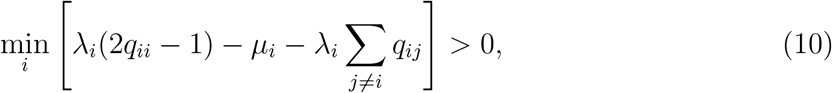

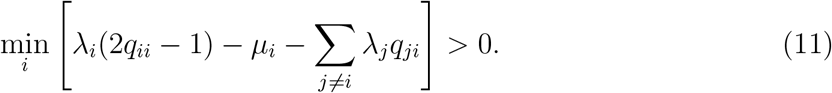

As both of these need to be positive, we can find the minimum of these, which will provide sufficient condition to escape MIE (see also Supplementary Information 1.4).

### 2.5 Predicting sensitivity to MIE

Given a fixed birth- and death-rate, can we predict at what missegregation rate a population will go extinct? Here we derive critical curves that separate viable from non-viable populations as a function of their turnover-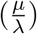 and missegregation rates (*β*). Herein we make three assumptions: (i) all missegregation events are possible (e.g. if parent after synthesis phase has 2*i* copies, then a daughter cell can have any integer in the range [0, 2*i*]); (ii) homogeneous turnover- and missegregation rates regardless of karyotype; and (iii) that the interval of viable karyotypes [*k*, *K*] is finite. Hereby we consider two types of viable karyotype intervals with different biological interpretations: intervals modeling the copy number of a single individual chromosome (e.g. *k* = 1, *K* = 5) and intervals modeling the ploidy of a cell (e.g. *k* = 22, *K* = 88). The former assumes there exists at least one single critical chromosome for which copy number must stay within a defined range for a cell to be viable. The latter treats all chromosomes as equal and models ploidy as the critical quantity.

To calculate the critical curves conditional on these assumptions, we consider the time evolution of the system given by the matrix form of eq. (8):

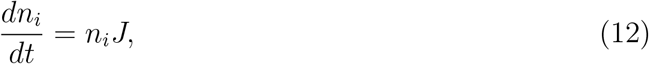

with J defined in eq. (9) and the row vector *n_i_* is the number of cells with copy number state i. It is clear that the system reaches a steady state when *n_i_J* = 0. Nontrivial solutions (i.e. those with nonzero *n_i_*) can be found by choosing functions for the mis-segregation rate *β* and death rate *μ* (which parameterise the matrix *J*) such that the dominant eigenvalue is zero (Supplementary Information 1.1). We also simulated the ODE given by eq. (8) until the karyotype distribution reached a steady state (Supplementary Fig. 1A,C,E). Numerical simulations confirmed that the theoretical critical curves separate exponential growth from population extinction (Supplementary Fig. 2).

We compared scenarios where the viable interval for ploidy is finite to scenarios where the viable interval for the copy number of individual chromosomes is finite (Fig. 2A). The latter contracted the viability region considerably more than the former, suggesting ploidy does not constrain the mis-segregation rates of cancer cell populations. When modeling single individual chromosomes, MIE was impossible at low turnover rates, even for very high *β* (Fig. 2B). This was because, as *β* → 1, none of the sister chromatids are properly segregated, i.e. they end up in the same cell, resulting in a high representation of cells with an even number of chromosomes. These in turn have a high enough fraction of viable daughter cells, sufficient to keep net growth above 0. In contrast, having more than one chromosome with finite viable karyotype intervals substantially contracted the viability region (Fig. 2B), albeit with diminishing costs in viability for each extra chromosome (Supplementary Fig. 4).

**Figure 2:**
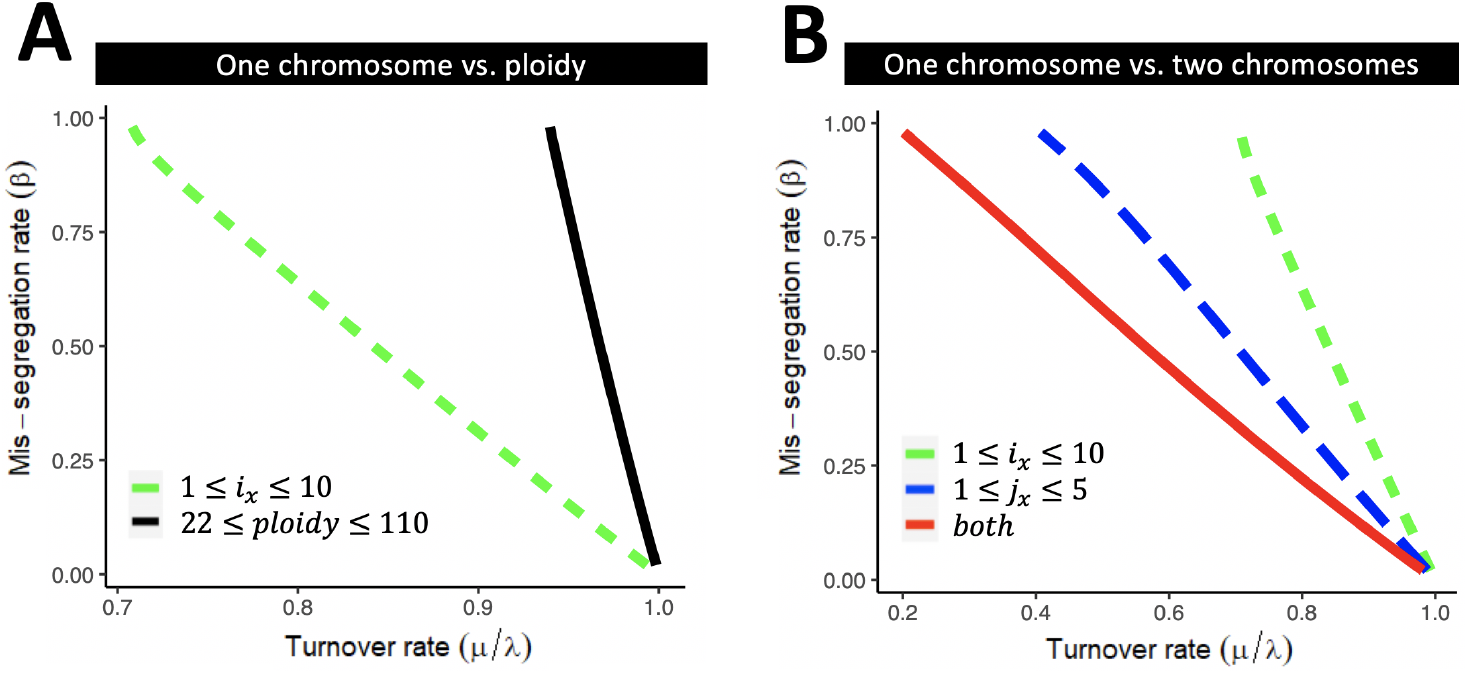
Predicting MIE as a function of homogeneous missegregation and turnover rates. (A-B) Critical curves were obtained by finding 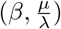 for which the maximum eigenvalues of the Jacobian (eq. (9)) is 0. (A) We consider two types of viable karyotype intervals with different biological interpretations: intervals modeling the copy number of a single individual chromosome (dashed line) and intervals modeling the ploidy of a cell (solid line). (B) We assume existence of two critical chromosomes *i* and *j*, with intervals of viable karyotypes [*k_i_*, *K_i_*] and [*k_j_*, *K_j_*] respectively. We calculate the critical curves assuming cell viability is restricted by only one of the two chromosomes (dashed lines), or by both chromosomes jointly (solid line). Note that the size of the Jaccobian is a function of 1 + *K* – *k*.

### 2.6 Quantification of mis-segregation and turnover rates across cancers

When measuring missegregation- and turnover rates in cancers we would expect these rates to lie below the predicted critical curves. To test this we quantified turnover- and missegregation rates at cellular resolution using scRNA-seq data from 15,464 single cells from the TISCH database [37]. Cells originated from 14 tumor biopsies across 12 patients spanning four cancer types across three tissue sites (Lung, Breast and Skin). We leverage the relation between turnover rates of cancers and their respective tissue site of origin [38, 39] (Methods 4.2.3), in order to learn to estimate turnover rate from transcriptomic signatures. A cell’s transcriptome is a channel of information propagation; it is a snapshot of how a cell interacts with and responds to its environment. Transcriptomic signatures have been used to infer various aspects about a tumor, ranging from the level of hypoxia [40], to its cell of origin [41], its mitotic index [42] and other surrogates of cell fitness and risk of disease progression [43, 44]. Cells co-existing in the same tumor, or in the same cell line [45, 46], often differ in their transcriptomes. Together these intra-tumor differences as well as inter-tumor differences in gene expression have the potential to inform how cells and tumors differ in their turnover rates.

After preprocessing (Methods 4.2.1), we performed Gene Set Variation Analysis (GSVA) [48] to quantify the expression activity of 1,629 REACTOME pathways [49] at single cell resolution. For each pathway involved in cell death and apoptosis (12 pathways), we calculated the median expression for a given tissue site and compared it to the median turnover rates [50–59] reported for cancers from the corresponding tissue site (Supplementary Table 1). Of the 12 tested pathways, five had an association with turnover rate (adjusted *R*^2^ ≥ 0.8; Methods 4.2.3), including “*FOXO-mediated transcription of cell death genes*” (Fig. 3A, adjusted *R*^2^ = 0.999; *P* = 0.07). We used this pathway signature to estimate turnover rate at single-cell resolution across the 14 tumors (Fig. 3B). All but one tumor had predicted turnover rates that were high, but below one (Supplementary Fig. 6), consistent with an expanding tumor mass. We note one exception, wherein a pre-treatment breast cancer sample had a median inferred turnover rate of 1.04 (Supplementary Fig. 6) – this case was excluded from further analysis.

**Figure 3:**
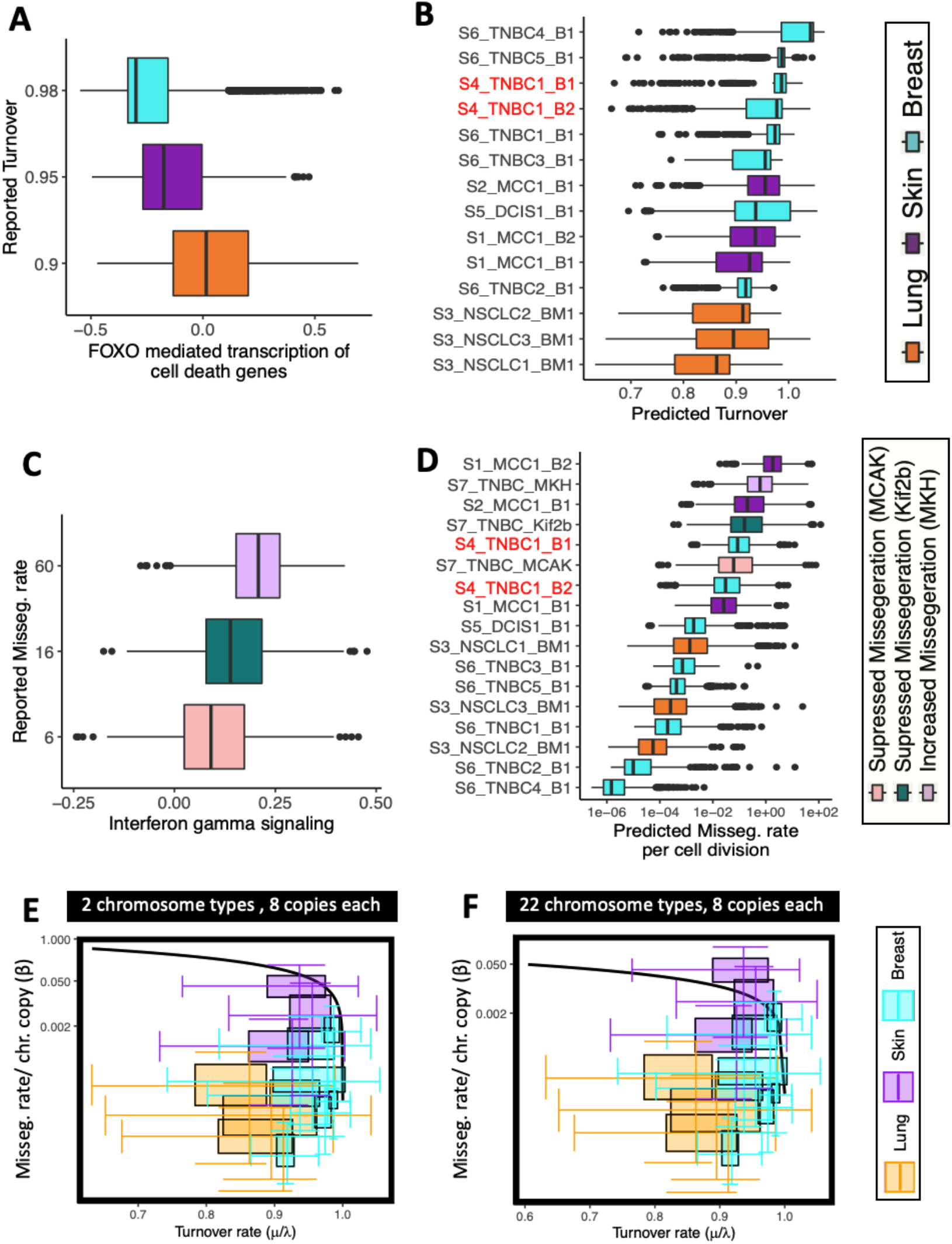
Predicting proximity of a PAN-cancer cohort to MIE. (A-D) Quantification of missegregation and turnover rates. (A) Expression of regulators of cell death genes (x-axis) varies across tissue sites (color code) along with turnover rates reported for tumors of the same origin (y-axis; adjusted *R*^2^ = 0.999; *P* = 0.07). (B) A regression model was built from (A) and used to predict turnover rate in 15,464 cells across 14 tumors. (C) Interferon Gamma gene expression (x-axis) measured in 7,879 cells from three human breast cancer cell lines [47] (color code) varies with their % lagging chromosomes quantified from imaging (y-axis; adjusted *R*^2^ = 0.88; *P* = 0.157). (D) A regression model was build from (C) and used to predict turnover rate in 23,343 cells across 17 tumors. (E,F) Missegregation-(*β*), and turnover rates (*μ*/λ) derived for 22,433 cells from 13 tumors across three tissue types (colors) are displayed alongside the critical curve predicted for two chromosome types (E) and all 22 autosomes (F). The interval of viable karyotypes is between one and eight copies for each chromosome type.

Given appropriate ground truth data, the same principle can be applied to estimate missegregation rates. To estimate mis-segregation rates at cellular resolution we used *Interferon Gamma Signaling* as a surrogate measure of chromosome missegregarions [47] (Methods 4.2.4). The rationale for this is that chromosome missegregations can trigger the formation of micronuclei. When micronuclei rupture, their genomic DNA spills into the cytosol. Cytosolic dsDNA is sensed by the cGAS–STING pathway [60], leading to induction of type I interferon stimulated genes [61, 62]. Missegregations lead to the upregulation of interferon production, which in turn subverts lethal epithelial responses to cytosolic DNA. To go from Interferon Gamma expression to mis-segregation rate we integrated aforementioned scRNA-seq dataset with 7,879 transcriptomes sequenced in Bakhoum et al. [47] (Methods 4.2.1). These transcriptomes originated from three cell lines, where members of the kinesin superfamily of proteins were knocked down to increase or decrease mis-segregation rate in a controlled fashion [47]. Live cell imaging of these cells to quantify the resulting missegregation rate was also available [47], allowing for a linear regression model to be fit on this data. As previously reported [47], Interferon Gamma Signaling was correlated to the % lagging chromosomes derived from imaging (Fig. 3C; adjusted *R*^2^ = 0.88; P = 0.157). The resulting model translates Interferon Gamma expression into units of mis-segregation rate per cell division and was used to estimate mis-segregation rates in the remaining scRNA-seq samples (Fig. 3D).

The number of chromosomes a mitotic cell has to segregate among daughter cells varies with ploidy. Therefore, the risk of mis-segregating at least one chromosome should increase with ploidy, rendering the per chromosome missegregation rate a quantity of interest. Calculating the missegregation rate per chromosome requires knowing the karyotype of each cell. To extract this information from the scRNA-seq data we extended an approach we previously described [43] to distinguish chromosome-arms affected by SCNAs from those that are copy number neutral (Methods 4.2.2). The resulting profiles were then clustered into subpopulations of cells with unique karyotypes as previously described [43] (Supplementary Fig. 5A), allowing for inference of mis-segregation rates per cell division per chromosome for each subpopulation (Supplementary Fig. 5B).

The variability in missegregation rates and the proximity of turnover rates to homeostasis warrants further investigation into whether increasing missegregation rate is a potential mechanism of extinction in these tumors. We therefore compared missegregation- and turnover rates derived from scRNA-seq data (Supplementary Fig. 6) to the critical curves. Since most tumors had high turnover rates, we focused on the critical curves at 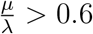 (Fig. 3E,F). Of note is the close proximity of the measured rates to the theoretical MIE curves. When imposing between one and eight copies on only two chromosome types, all tumors had median missegregation- and turnover rates that were compatible with our viability predictions (Fig. 3E). When imposing between one and eight copies on all 22 autosomes, one of the three skin cancers slipped into the region predicted as non-viable. But for all remaining tumors, even when considering all 22 autosomes, the majority of cells stayed in the viable region (Fig. 3F).

### 2.7 Intra-tumor heterogeneity in mis-segregations and turnover

The critical curves shown in Fig. 3E,F assume a cell population with homogeneous missegregation and turnover rates. The scRNA-seq derived results however suggest that both are likely heterogeneous in reality. We therefore asked whether relaxing this assumption changes the critical curves. To model intra-tumor heterogeneity in missegregation rates, we looked at their relation to ploidy and chromosome copy number. While no significant association between ploidy and turnover rate was evident, the relationship between ploidy and missegregation rate per chromosome per cell division showed a surprising resemblance to the recently hypothesized fitness function of ploidy [63] (Supplementary Figs. 7, 6B,C). Hereby the commonly observed near-triploid karyotype [35, 64, 65] stands out as a local maximum. A sinus function was therefore chosen to model mis-segregation as a function of ploidy (adjusted *R*^2^ = 0.80; Estimated Variance: 49%, Supplementary Fig. 7D). A linear association between copy number and missegregation rate was also observed for three of the 22 individual autosomes (adjusted *R*^2^ > 0.1; *P* << 1*E* – 5, Supplementary Fig. 7A-C). The observed relation between missegregation rate per chromosome and ploidy (either of specific chromosomes or in aggregate), is an opportunity to model missegregation rate as a function of ploidy, thereby accounting for intra-tumor heterogeneity.

Modeling missegregation rate as linear or sinusoidal functions of the copy number of chromosome 4 and overall ploidy respectively (Fig. 4A,B), we calculated how parameters of both functions shape the critical curve (Fig. 4C,D; Supplementary Fig. 1B,D,F). In both cases the population evolved toward the karyotype with the lowest mis-segregation rate (Fig. 4E,F). Unless turnover rates are exceedingly high, this convergence to the global minimum mis-segregation rates effectively rendered MIE impossible (Fig. 4G,H). The absolute value of that minimum explains the difference in the location of the critical curve between the two scenarios (Fig. 4G,H), and the overall low risk of MIE in large cell populations.

**Figure 4:**
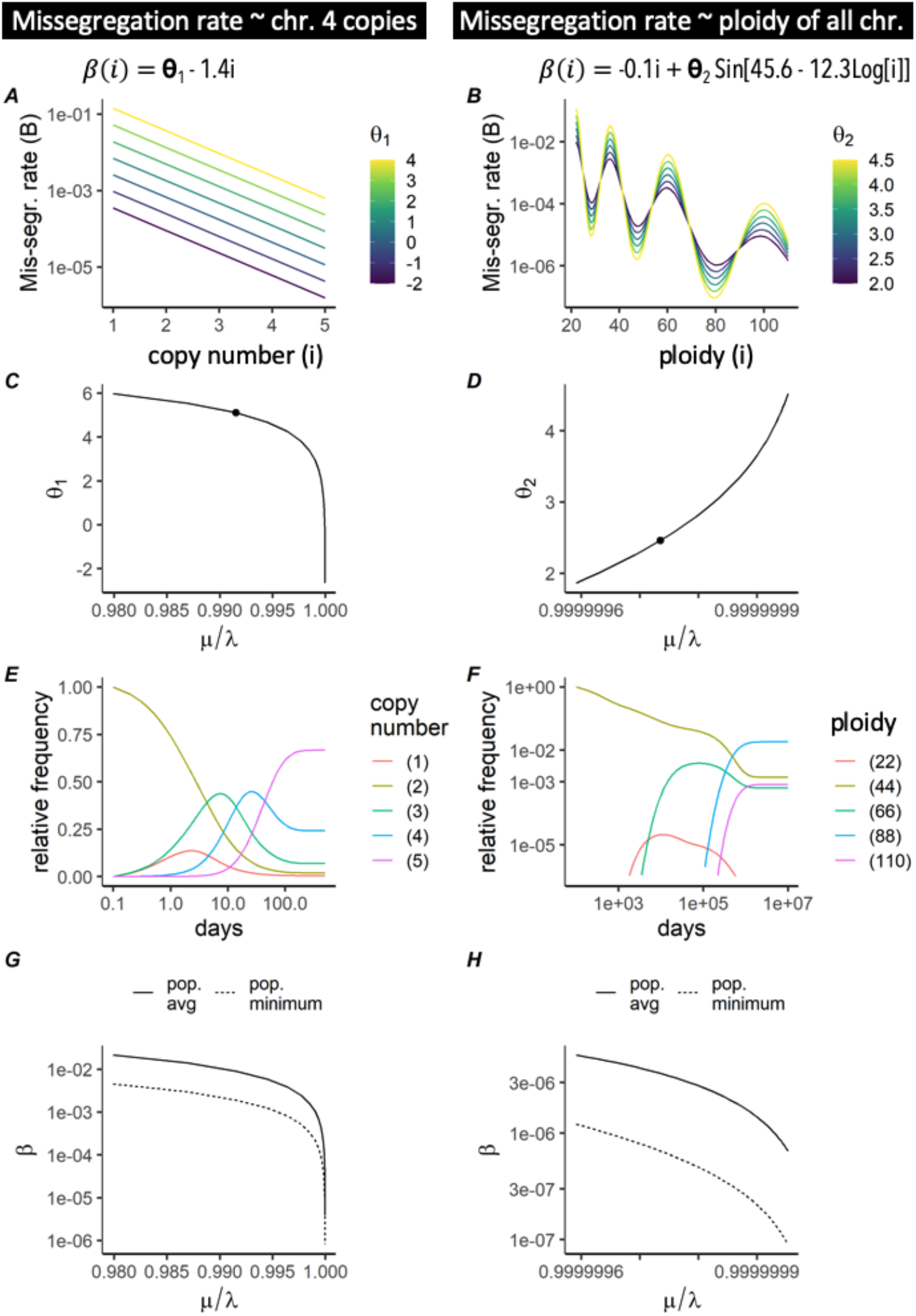
Predicting MIE when missegregation rates are heterogeneous. (A,B) Missegregation rate is modeled as a function of copy number or ploidy (x-axis), with color coded shape parameters *θ*_1_ (A) and *θ*_2_ (B). We consider the copy number of either chromosome 4 alone (A) or of all 22 autosomes in aggregate, i.e. ploidy (B). Varying *θ*_1_, *θ*_2_ yields different missegregation rates (*β*). (C) Critical curve was obtained for (A) by finding 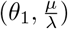 for which the maximum eigenvalues of the Jacobian (eq. (9)) is 0. (D) Critical curve was obtained for (B) in the same manner as in (C), but here equations were solved for 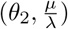. (E,F) We used the parameters highlighted in (C,D) to simulate missegregations until the karyotype composition reached a steady state. (G,H) The eigenvectors corresponding to the eigenvalues found in C/D are the steady state karyotype proportions. These are used in conjunction with the kernels in A/B to determine the population average and minimum missegregation rates.

If we also model heterogeneous death rates, the interplay between mis-segregation and death rate, rather than the minimum mis-segregation rate alone, determine whether MIE will occur. More generally, when both missegregation- and death rates are heterogeneous, a necessary condition for MIE is that karyotypes with low mis-segregation rates must also have high death rates (Supplementary Fig. 3A,B). This condition is exactly identical to the sufficient condition for ruling out MIE described by eq. (10), and was corroborated by combinations of kernels of missegregation- and death rate (Supplementary Fig. 3C). Taken together these results suggests that heterogeneous missegregation rates can protect a population from extinction.

## 3 Discussion

We have presented a general approach for modeling whole chromosome missegregations, including a deterministic mathematical framework and scRNA-seq analysis methods to infer effective model parameters. In contrast to prior models of chromosome missegregations [32–34], our model does not rest on the assumption that the fitness effect of a mis-segregation is the same, regardless of the karyotype context in which it happens. This feature offers the flexibility to identify potential synergies between copy number changes of multiple chromosomes [66]. A second difference to prior models of missegregations [32], is the decoupling of a cell’s life cycle from the life cycle of individual chromosomes. This allows simulating intratumor heterogeneity in mis-segregation, death- and proliferation rates across cells, which can manifest as temporal variations in these rates (when the karyotype composition changes over time).

Theoretical analysis of the mathematical model has shown the existence of a potential mechanism of tumor control through the region in parameter space we have called MIE. As a first application of the model we have thus focused on the identification of critical curves that separate viable populations from MIE, as a function of their turnover- and missegregation rates. A central assumption of these calculations is that cells are not viable unless they carry a certain number of copies of a given chromosome, that must lie within a predefined interval. To our knowledge, Fig. 2 contains the first predictions of MIE that consider viable karyotype intervals of multiple chromosomes simultaneously, as well as the turnover rate of the population. We compared these theoretical critical curves to missegregation- and turnover rates inferred from scRNA-seq of 13 tumors across four cancer types from three tissue sites. The majority of tumors across all tissue sites studied had missegregation- and turnover rates that were compatible with our viability predictions (Fig. 3E,F). This remained true when a dependency of missegregation rates on ploidy was introduced. In populations with heterogeneous mis-segregation rates, the subpopulation with the minimum mis-segregation rate protects the population from extinction (Fig. 4G,H). Our results emphasize that large, heterogeneous tumors have an inbuilt protection from MIE. That each tumor consists of cells with heterogeneous missegregation rates, the measurement being just the population-average rate, is a likely scenario supported by recent results [43, 67]. Karyotypes associated with low missegregation rates act as a stabilizing refuge, protecting the population from extinction. Intra-tumor heterogeneity, including heterogeneity in missegregation rates, increases as tumors progress. Our predictions suggest that this intra-tumor heterogeneity renders MIE unlikely.

The model raises some important theoretical questions related to malignant and non-malignant cells. In particular, it is well known that normal cells maintain a level of homeostasis through a balanced turnover rate *μ*/λ ≈ 1. This seems to imply that *all* normal cells lay at the MIE boundary and are even more sensitive to MIE than malignant cells. Any finite missegregation rate would thus lead to the slow removal of normal cells over time. There are two potential explanations for this behavior. A likely explanation is given by our assumption that the birth rate is independent of population size. It is easy to see that introducing dependency on the total populations size (e.g. carrying capacity) could alleviate this issue as cell death would increase the birth rate in order to return to homeostasis. An alternative explanation is that this is just another natural aging mechanism through which normal cells are slowly displaced. Transformed cells often lose the homeostatic control mechanisms and so are likely less susceptible to contact inhibition.

Limitations of our approach include unknown precision of mis-segregation and turnover rates inferred from scRNA-seq. In line with prior reports [68], mis-segregation rates were higher in higher stage cancer, while normal tissue used as control had the lowest missegregation rates (Supplementary Fig. 9). The number of subpopulations with distinct karyotypes was also by trend higher in late stage tumors – a finding that is also consistent with prior reports [68–70]. Other limitations include those of ordinary differential equation models, such as the lack of stochasticity, rendering all conclusions valid only for large populations, where all karyotypes are accessible and represented. Understanding how missegregations shape extinction events in early stage cancers would require a different approach. Our model does not explicitly account for several biological mechanisms which are relevant to karyotype evolution, including WGD, missegregation induced apoptosis in the subsequent G1 phase of the cell cycle, and the formation of micronuclei. Extensions to model these phenomena are discussed in Supplementary Information 1.5.

Future applications of this model will include studying the fitness costs and benefits of high ploidy. Coexistence of cancer cells at opposite extremes of the ploidy spectrum occurs frequently in cancer and missegregations are a major contributor to heterogeneous ploidy states within a population. Our model can help understand how much robustness high ploidy confers to the sudden genome-dosage imbalance caused by a missegregation event [13] and can help quantify the energetic requirements of high ploidy cells. Modeling intra-tumor heterogeneity in mis-segregation rates and their effect on karyotype evolution over hundreds of generations can reveal how selection acts upon coexisting karyotypes, as a powerful tool for genotype-to-phenotype mapping in various microenvironments.

## 4 Materials and Methods

### 4.1 Numerical Simulations

Numerical simulations were performed for a range of input parameters (*β*, *μ*) to validate the predicted critical curves. All simulations had a uniform diploid population as initial condition. Each simulation ran until a quasi-steady-state (QSS) had been reached, considered to occur when the rate of change in karyotype composition was less than 0.1%*day*^-1^ (although the cell population may still be growing or shrinking – therefore “quasi”). Upon satisfaction of this condition, simulations with a positive rate of change in the total cell population were considered to be in the exponential growth regime.

In order to determine the population average missegregation rates *β_pop_* and death rates *μ_pop_* which are viable QSS’s for the system, we performed numerical simulations for each pairwise combination of missegregation- and death rate kernels – *β*(*i*, *B*) and *μ*(*i*, *M*) respectively. For each pairwise combination of kernels, simulations were performed using large, manually curated ranges for the input parameters (*B*, *M*), before *β_pop_* and *μ_pop_* were calculated based on the QSS reached by the system. Population average missegregation rate (*β_pop_*) was defined as fraction of divisions in which a missegregation occurs (i.e. 1 – (1 – *β*)^*i*^).

Numerical simulations were performed using R, all code is available on Github.

### 4.2 Quantification of ploidy, missegregation- and turnover rates across cancers

To parametrize our ODE, we derive ploidy, mis-segregation and turnover rates from an integrated scRNA-seq dataset. Fourteen samples from 12 patients across four cancer types were downloaded from the TISCH database [37] and analyzed as follows.

#### 4.2.1 scRNA-Seq data integration

Filtered gene-barcodes matrices containing only barcodes with UMI counts passing threshold for cell detection were imported to Seurat v4.0 for downstream analysis. Barcodes with fewer than 500 genes expressed or more than 25% mitochondrial UMIs were filtered out; genes expressed in fewer than 3 barcodes were also excluded. Raw counts of different datasets were merged using merge function. Standard library size and log-normalization was performed on raw UMI counts using NormalizeData, and top 5000 most variable genes were identified by the “vst” method in FindVariableFeatures. S and G2/M cell cycle phase scores were assigned to cells based on previously defined gene sets(9) using CellCycleScoring function. Normalized UMI counts were further scaled using ScaleData function by regressing against total reads count, % of mitochondrial UMIs, and cell cycle phase scores (G2M.Score, and S.Score) to mitigate the effects of sequencing depth and cell cycle heterogeneity. UMAP analysis of the data after these normalization steps were performed shows that cells cluster by cell types, rather than by study, which indicated that the batch effects were delimited by normalization and scaling (Supplementary Fig. 8). Because 10X measures UMI (copy of transcripts), not number of reads mapped to genes and because GSVA uses the ranking of the transcripts, the data from different experiments are comparable when quantified into pathway activity.

#### 4.2.2 Estimating ploidy from scRNA-Seq

Our goal was to distinguish chromosome(-arm)s affected by SCNAs from those that are copy number neutral, given a set of tumor cells and normal cells from same patient. Normal cells (often immune cells) were not of the same type as tumor cells (epithelial). Hence, using them directly as a control to calculate absolute copy number in tumor cells is problematic: immune cells express different numbers of genes (often less), and may have a different viability during scRNA-seq library preparation. To overcome this challenge, we assume that at least one chromosome is diploid in all tumor cells and that most SCNAs are clonal (i.e. they affect all tumor cells) [71].

We first sort chromosomes by the p-value of differential chromosome-specific gene expression between tumor and normal cells in descending order of significance. We chose an *x* ∈ {1..22} and define 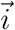 and 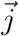 as the vectors of the first *x* and last (22–*x*) chromosomes in the sorted set respectively. We then proceed as follows: (i) We assume all chromosomes in 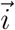 have identical (diploid) copy number in tumor and normal cells. The average ratio of expression between tumor and normal cells for these *x* chromosomes should thus be 1. Deviation from 1 is the bias (*ϵ_x_*) we estimate between tumor and normal cells: 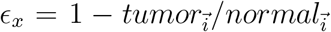. (ii) We calculate the vector 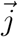 of copy numbers of chromosomes with SCNAs (all except the first *x*), with entries *j_k_* as: *j_k_* = (2/*ϵ_x_*) * (*tumor_k_/normal_k_*) for each chromosome *k*. (iii) We evaluate deviation of 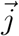 from the closest integers: 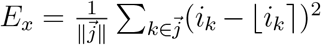. Repeating steps (i-iii) for all possible values of *x* lets us choose the *x*\*:= arg min_*x*_ *E_x_* (Supplementary Fig. 10). This classifies the last (22 – *x*\*) chromosomes as chromosomes affected by SCNAs and gives us their absolute copy numbers in the respective 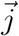.

We then used LIAYSON [43] to classify cells into subpopulations with distinct karyotypes. The number of subpopulations was by trend higher in tumors of high ploidy (Spearman *r* = 0.484; *P* = 0.079), but lower in tumors with high turnover rates (Spearman *r* = −0.489; *P* = 0.076; Supplementary Fig. 6). We also observed that a clone’s ploidy was positively associated with its variance in turnover rates (Spearman *r* = 0.41; *P* = 7.9*E* – 5). This positive association was also observed when considering each of the three tissue sites (Breast, Lung, Skin) individually, albeit it only reached significance in Lung cancer (Spearman *r* = 0.63; *P* = 8.1*E* – 6).

#### 4.2.3 Estimating turnover rates from scRNA-seq

Reported proliferation rates from tumors correlate to turnover rates from their respective normal tissue of origin (Pearson *r* = 0.93, *P* = 0.021; Supplementary Table 1). The same is true for reported cancer cell death rates, which also correlate to the death rates of their tissue of origin (Pearson *r* = 0.92, *P* = 0.025; Supplementary Table 1). The relation between turnover rates of cancers and their respective tissue site of origin [38, 39], is an opportunity to learn how to read these rates from transcriptomic signatures. We performed Gene Set Variation Analysis (GSVA) [48] to quantify the expression activity of 1,629 REACTOME pathways [49] in a cumulative total of 43,596 single cells from 15 samples across three tissue sites. For each pathway involved in cell death and apoptosis (12 pathways), we calculated the average expression among all cells of a given tissue site and used it to model the median turnover rates [50–59] reported for cancers from the corresponding tissue site (Supplementary Table 1):

We fitted a linear regression model on the combined dataset as follows:

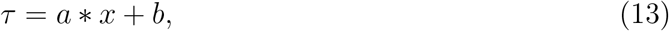

where *x* is the average pathway expression signature per cancer and *τ* is the turnover rate reported in literature for that cancer type. Of all tested pathways, five had an association with turnover rate (adjusted *R*^2^ ≥ 0.8), including “*FOXO-mediated transcription of cell death genes*” (adjusted *R*^2^ = 0.999; *P* = 0.07). This pathway signature was then used to estimate *τ* in each single cell across the four cancer types (Fig. 3B). We set birth rate to 1, and used *μ*:= *τ* as death rate for all further mathematical modeling.

#### 4.2.4 Estimating missegregation rate from scRNA-seq

Interferon Signaling has been proposed as potential surrogate measure for CIN [47]. To predict missegregation rate from expression of genes involved in *Interferon Gamma Signaling*, we used a similar approach as for turnover rate. We fitted a linear regression on the breast cancer data from [47] as follows:

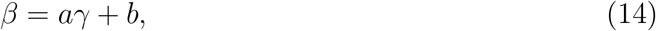

where *β* is the log2 of observed percentage of cells with lagging chromosomes and *γ* the Interferon Gamma Signaling activity as quantified with GSVA (adjusted R-square=0.999, p-value=0.0103; Fig. 3C). The resulting model was then used to predict missegregation rate in 15,464 single cells from the 14 tumor samples (Fig. 3D). We divided the predicted missegregation rate by ploidy to obtain *β* for all further mathematical modeling.

The hereby obtained relationship between missegregation rate and karyotype (Supplementary Fig. 7D), was similar to how karyotype and fitness are thought to be linked [63]. Namely, triploid karyotypes had higher mis-segregation rates and missegregation rates of euploid states tended to decrease with ploidy. This trend was only evident when looking at the ploidy spectrum across all four cancer types. Variability in ploidy was too low to test if this observation holds across tumors of a given type and especially across subpopulations within a given tumor. That ploidy and cancer type are confounded prevents any causal conclusions to be drawn from this analysis. This correlation is however to be expected, because ploidy is highly cancer types specific [72–74].

## 5 Acknowledgements

This work was supported by NCI grants R00CA215256 and 1R01CA266727-01A1 awarded to N. Andor. P.M. Altrock and N. Andor acknowledge support through the NCI, part of the NIH, under grant number P30-CA076292 and from the Moffitt Cancer Center Evolutionary Therapy Center of Excellence. P.M. Altrock also acknowledged funding from Richard O. Jacobson Foundation, and the William G. ‘Bill’ Bankhead Jr and David Coley Cancer Research Program (20B06). The funders had no role in study design, data collection and analysis, decision to publish, or preparation of the manuscript.

## 6 Code availability

The full general discrete model presented in section 2.2 is implemented in R, and is available at https://github.com/MathOnco/MIEmodel.

## 7 Competing interests

The authors declare no competing interest.

## Supplementary Information: Legends

Supplementary Table 1: ***In vivo* tumor dynamics across nine cancer types**. Birth rates taken from [50] and growth rates taken from multiple sources (cited in third column). The death rate is inferred. Many of the birth rates are comparable, but the net growth rates vary over orders of magnitude, which implies very different death rates. Last column shows percent contribution of respective normal human cell types to a total daily mass turnover of 80 g (taken from [75]) – these are correlated to both birth and death rates reported in tumors (Pearson *r* ≥ 0.923; *P* < 0.026).

Supplementary Figure 1: **Predicted changes in missegregation rate during tumor evolution.** (A-B) Critical curves displayed as a function of the population average missegregation rate (y-axis) and departure from homeostasis (1 - turnover rate; x-axis). λ:= 1 in all calculations, i.e. cells divide once per day. Copy number ranges from [1, 8] and death rate is independent of copy number (*μ*(*i*):= *M*), whereas missegregation is either also independent (A, *β*(*i*):= Θ_1_) or dependent on copy number (B, *β*(*i*):= Θ_2_ – 1.4*i*). (C-D) Population average missegregation and turnover rates over time are shown for three parameter combinations *θ* ∈ *a*, *b*, *c* highlighted in (A,B) respectively. (E-F) Evolution of karyotype composition for each of the three parameter combinations is also shown.

Supplementary Figure 2: **Numerical simulations confirm theoretical MIE curves**. Numerical simulations confirm that the theoretical critical curves shown in Fig. 2A,B separate exponential growth from population extinction.

Supplementary Figure 3: **Heterogeneous missegregation and death rates can render MIE impossible**. Kernels used to model intra-tumor heterogeneity in missegregation-(A) and turnover rates (B). (C) Critical curves calculated assuming homogeneous or heterogeneous missegregation and turnover rates are compared with each other. Functions used to model heterogeneous rates are displayed as row and column labels and have ploidy (*i*) as parameter. For each of these functions, we calculated the critical missegregation rate parameter (*B*) and death rate parameter (*M*). Note that for *M* = 1, the turnover rate function in the first column becomes the constant rate from the second column. MIE requires *M* → 1, explaining why critical curves look identical in both columns. (D) We also simulated the ODE assuming combinations of the functional forms of missegregation and turnover rates shown in C. Shown here is the number of days needed to reach steady state karyotype composition as a function of population average missegregation rate.

Supplementary Figure 4: **Risk of MIE as a function of the number of chromosome types with finite viable copy number intervals**. (A-C) Critical curves were obtained by finding 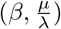 for which the maximum eigenvalues of the Jacobian (eq. (9)) is 0. We assume existence of up to *m* = 22 critical chromosomes, where intervals of viable karyotypes for each chromosome type *x* ∈ 1..*m* are defined by *k_x_* ≤ *i_x_* ≤ *K_x_*. We calculate the critical curves assuming cell viability is restricted by all 22 of the chromosomes or by only a subset of them (color code). Hereby we assume each chromosome must have at least one and no more than three (A), five (B) or eight copies respectively (C).

Supplementary Figure 5: **Karyotype profiles inferred for two metastatic TNBC samples**. (A) Copy number profiles from two of the metastatic samples from patient *S4_TNBC1* (marked red in Fig. 3B,D) were clustered into subpopulations of cells with unique karyotypes (left color code). The two metastatic samples originate from the same patient and have significant similarities in their karyotype profiles, even though they were called independently from each other. (B) Inferred mis-segregation rates per cell division per chromosome for each subpopulation in (A).

Supplementary Figure 6: **Variability in turnover, missegregation and ploidy across coexisting subpopulations in 14 tumors**. Every column represents a subpopulation of cells with unique karyotype. Subpopulations from the same tumor are grouped together. Their ploidy (A), departure from homeostasis (B) and log2-transformed % missegregation rate (C) are shown.

Supplementary Figure 7: **Relationship between karyotype and missegregation rates**. Missegregation rates per chromosome (y-axis) vary with the copy number (x-axis) of specific chromosomes (A-C) and with ploidy (i.e. copy numbers of all chromosomes in aggregate; D).

Supplementary Figure 8: **Effect of normalization and scaling for integration of scRNA-seq datasets.** (A-C) UMAP plots using the normalized and scaled data, with the cells labeled by (A) sample name, (B) cancer type of the patients and (C) cell types. Tumor cells cluster by sample origin, and tumor cells from similar cancer types are closer to each other. The PBMC sample of the MCC patient clusters with other normal cells, instead of with the tumor cells from the same patient. Normal cells cluster mostly by cell type. (D-F) Analogous plots to (A-C), but generated using non-normalized data. Generally the cells are more clustered by study, and the tumor cells from different samples are less distinguished, emphasizing the need for normalization and scaling.

Supplementary Figure 9: **Mis-segregation rates inferred from scRNA-seq derived gene signatures**. Distribution of median mis-segregation rate across all sequenced G1 cells of a given sample are grouped by disease stage (A) or site (B). PBMC:= peripheral blood mononuclear cells.

Supplementary Figure 10: **Classification of aneuploid chromosomes with scRNA-Seq derived bias profiles**. Minimum deviation from integer copy numbers (y-axis) guides the choice of x (highlighted red). The more to the right the minimum, the more chromosomes are assumed to be diploid. Shown are the bias profiles for 15 scRNA-seq samples, sorted by global minimum. Samples from late-stage tumors (stage IV) have by trend more non-diploid chromosome arms and their global minimum is higher. Range of x-axis varies because only chromosomes that express a sufficient number of genes in both tumor and normal cells were considered.

## Notes

### Competing Interest Statement

The authors have declared no competing interest.

### Summary of Updates

We have analyzed scRNA-seq data to quantify intra-tumor heterogeneity in mis-segregation rates and in karyotypes and used these as effective parameters of our mathematical model. This allowed us to predict missegregation-induced extinction in context of a more realistic manifestation of intra-tumor heterogeneity (in both mis-segregation rates and the resulting karyotypes). An important conclusion from this additional analysis was that, in a population with heterogeneous mis-segregation rates, the cell subpopulation with the minimum mis-segregation rate explains differences in extinction likelihood, suggesting heterogeneous missegregation rates can protect a population from extinction.

## References

[1] Qinghua Shi and Randall W. King. Chromosome nondisjunction yields tetraploid rather than aneuploid cells in human cell lines. Nature, 437(7061):1038–1042, October 2005.

[2] Sarah L. Thompson and Duane A. Compton. Examining the link between chromosomal instability and aneuploidy in human cells. The Journal of Cell Biology, 180(4):665–672, February 2008.

[3] D Choma, JP Daures, X Quantin, and JL Pujol. Aneuploidy and prognosis of nonsmall-cell lung cancer: a meta-analysis of published data. British journal of cancer, 85(1):14–22, 2001. Publisher: Nature Publishing Group.

[4] Mallika Siva Donepudi, Kasturi Kondapalli, Seelam Jeevan Amos, Pavithra Venkanteshan, and others. Breast cancer statistics and markers. Journal of cancer research and therapeutics, 10(3):506, 2014. Publisher: Medknow Publications.

[5] Axel Walther, Richard Houlston, and Ian Tomlinson. Association between chromosomal instability and prognosis in colorectal cancer: a meta-analysis. Gut, 57(7):941–950, 2008. Publisher: BMJ Publishing Group.

[6] Maximilian Lennartz, Sarah Minner, Sophie Brasch, Hilko Wittmann, Leonard Paterna, Katja Angermeier, Eray Öztürk, Rami Shihada, Mingu Ruge, Martina Kluth, and others. The combination of DNA ploidy status and PTEN/6q15 deletions provides strong and independent prognostic information in prostate cancer. Clinical Cancer Research, 22(11):2802–2811, 2016. Publisher: AACR.

[7] Daniel Stieber, Anna Golebiewska, Lisa Evers, Elizabeth Lenkiewicz, Nicolaas HC Brons, Nathalie Nicot, SÚbastien Bougnaud, Frank Hertel, Rolf Bjerkvig, Laurent Vallar, and others. Glioblastomas are composed of genetically divergent clones with distinct tumourigenic potential and variable stem cell-associated phenotypes. Acta neuropathologica, 127(2):203–219, 2014. Publisher: Springer.

[8] Samuel F. Bakhoum, Lilian Kabeche, John P. Murnane, Bassem I. Zaki, and Duane A. Compton. DNA-damage response during mitosis induces whole-chromosome missegregation. Cancer Discovery, 4(11):1281–1289, November 2014.

[9] Makoto T. Hayashi, Anthony J. Cesare, James A. J. Fitzpatrick, Eros Lazzerini-Denchi, and Jan Karlseder. A telomere-dependent DNA damage checkpoint induced by prolonged mitotic arrest. Nature Structural & Molecular Biology, 19(4):387–394, March 2012.

[10] Rune Troelsgaard Pedersen, Thomas Kruse, Jakob Nilsson, Vibe H. Oestergaard, and Michael Lisby. TopBP1 is required at mitosis to reduce transmission of DNA damage to G1 daughter cells. The Journal of Cell Biology, 210(4):565–582, August 2015.

[11] Sheroy Minocherhomji, Songmin Ying, Victoria A. Bjerregaard, Sara Bursomanno, Aiste Aleliunaite, Wei Wu, Hocine W. Mankouri, Huahao Shen, Ying Liu, and Ian D. Hickson. Replication stress activates DNA repair synthesis in mitosis. Nature, 528(7581):286–290, December 2015.

[12] Samuel F. Bakhoum, Lilian Kabeche, Duane A. Compton, Simon N. Powell, and Holger Bastians. Mitotic DNA Damage Response: At the Crossroads of Structural and Numerical Cancer Chromosome Instabilities. Trends in Cancer, 3(3):225–234, March 2017. Publisher: Elsevier.

[13] Jason M. Sheltzer, Julie H. Ko, John M. Replogle, Nicole C. Habibe Burgos, Erica S. Chung, Colleen M. Meehl, Nicole M. Sayles, Verena Passerini, Zuzana Storchova, and Angelika Amon. Single-chromosome Gains Commonly Function as Tumor Suppressors. Cancer Cell, 31(2):240–255, February 2017.

[14] Neil J Ganem, Susana A Godinho, and David Pellman. A mechanism linking extra centrosomes to chromosomal instability. Nature, 460(7252):278–282, 2009. Publisher: Nature Publishing Group.

[15] Norman Ertych, Ailine Stolz, Albrecht Stenzinger, Wilko Weichert, Silke Kaulfuß, Peter Burfeind, Achim Aigner, Linda Wordeman, and Holger Bastians. Increased microtubule assembly rates influence chromosomal instability in colorectal cancer cells. Nature Cell Biology, 16(8):779–791, August 2014.

[16] Samuel F. Bakhoum, Sarah L. Thompson, Amity L. Manning, and Duane A. Compton. Genome stability is ensured by temporal control of kinetochore-microtubule dynamics. Nature Cell Biology, 11(1):27–35, January 2009.

[17] Samuel F. Bakhoum and Duane A. Compton. Kinetochores and disease: keeping microtubule dynamics in check! Current Opinion in Cell Biology, 24(1):64–70, February 2012.

[18] Darcy Bates and Alan Eastman. Microtubule destabilising agents: far more than just antimitotic anticancer drugs. British Journal of Clinical Pharmacology, 83(2):255–268, February 2017.

[19] Paola Leopardi, Francesca Marcon, Gabriella Dobrowolny, Andrea Zijno, and Riccardo Crebelli. Influence of donor age on vinblastine-induced chromosome malsegregation in cultured peripheral lymphocytes. Mutagenesis, 17(1):83–88, January 2002.

[20] Hee-Sheung Lee, Nicholas C. O. Lee, Natalay Kouprina, Jung-Hyun Kim, Alex Kagansky, Susan Bates, Jane B. Trepel, Yves Pommier, Dan Sackett, and Vladimir Larionov. Effects of Anticancer Drugs on Chromosome Instability and New Clinical Implications for Tumor-Suppressing Therapies. Cancer Research, 76(4):902–911, February 2016.

[21] Samuel F. Bakhoum, Lilian Kabeche, Matthew D. Wood, Christopher D. Laucius, Dian Qu, Ashley M. Laughney, Gloria E. Reynolds, Raymond J. Louie, Joanna Phillips, Denise A. Chan, Bassem I. Zaki, John P. Murnane, Claudia Petritsch, and Duane A. Compton. Numerical chromosomal instability mediates susceptibility to radiation treatment. Nature Communications, 6:5990, January 2015.

[22] Chunyan Dai, Feifei Sun, Chunpeng Zhu, and Xun Hu. Tumor environmental factors glucose deprivation and lactic acidosis induce mitotic chromosomal instability–an implication in aneuploid human tumors. PLoS One, 8(5):e63054, 2013. Publisher: Public Library of Science.

[23] Miyako Kondoh, Noritaka Ohga, Kosuke Akiyama, Yasuhiro Hida, Nako Maishi, Alam Mohammad Towfik, Nobuo Inoue, Masanobu Shindoh, and Kyoko Hida. Hypoxia-induced reactive oxygen species cause chromosomal abnormalities in endothelial cells in the tumor microenvironment. PloS one, 8(11):e80349, 2013. Publisher: Public Library of Science.

[24] Sally M. Dewhurst, Nicholas McGranahan, Rebecca A. Burrell, Andrew J. Rowan, Eva Grönroos, David Endesfelder, Tejal Joshi, Dmitri Mouradov, Peter Gibbs, Robyn L. Ward, Nicholas J. Hawkins, Zoltan Szallasi, Oliver M. Sieber, and Charles Swanton. Tolerance of whole-genome doubling propagates chromosomal instability and accelerates cancer genome evolution. Cancer Discovery, 4(2):175–185, February 2014. 00068.

[25] Takeshi Fujiwara, Madhavi Bandi, Masayuki Nitta, Elena V. Ivanova, Roderick T. Bronson, and David Pellman. Cytokinesis failure generating tetraploids promotes tumorigenesis in p53-null cells. Nature, 437(7061):1043–1047, October 2005.

[26] Mario Vitale. Intratumor BRAFV600E heterogeneity and kinase inhibitors in the treatment of thyroid cancer: a call for participation. Thyroid: official journal of the American Thyroid Association, 23(4):517–519, April 2013. 00000.

[27] Samuel F. Bakhoum and Duane A. Compton. Chromosomal instability and cancer: a complex relationship with therapeutic potential. The Journal of Clinical Investigation, 122(4):1138–1143, April 2012.

[28] Sarah L. Thompson and Duane A. Compton. Proliferation of aneuploid human cells is limited by a p53-dependent mechanism. The Journal of Cell Biology, 188(3):369–381, February 2010.

[29] Stefano Santaguida, Amelia Richardson, Divya Ramalingam Iyer, Ons M’Saad, Lauren Zasadil, Kristin A. Knouse, Yao Liang Wong, Nicholas Rhind, Arshad Desai, and Angelika Amon. Chromosome mis-segregation generates cell cycle-arrested cells with complex karyotypes that are eliminated by the immune system. Developmental cell, 41(6):638–651.e5, June 2017.

[30] Rebecca Roylance, David Endesfelder, Patricia Gorman, Rebecca A Burrell, Jil Sander, Ian Tomlinson, Andrew M Hanby, Valerie Speirs, Andrea L Richardson, Nicolai J Birkbak, Aron C Eklund, Julian Downward, Maik Kschischo, Zoltan Szallasi, and Charles Swanton. Relationship of extreme chromosomal instability with long-term survival in a retrospective analysis of primary breast cancer. Cancer epidemiology, biomarkers & prevention: a publication of the American Association for Cancer Research, cosponsored by the American Society of Preventive Oncology, 20(10):2183–2194, October 2011. 00039 PMID: 21784954.

[31] Nicolai J Birkbak, Aron C Eklund, Qiyuan Li, Sarah E McClelland, David Endesfelder, Patrick Tan, Iain B Tan, Andrea L Richardson, Zoltan Szallasi, and Charles Swanton. Paradoxical relationship between chromosomal instability and survival outcome in cancer. Cancer research, 71(10):3447–3452, May 2011. 00068 PMID: 21270108.

[32] Y Gusev, V Kagansky, and W. C Dooley. A stochastic model of chromosome segregation errors with reference to cancer cells. Mathematical and Computer Modelling, 32(1):97–111, July 2000.

[33] Y. Gusev, V. Kagansky, and W. C. Dooley. Long-term dynamics of chromosomal instability in cancer: A transition probability model. Mathematical and Computer Modelling, 33(12):1253–1273, June 2001.

[34] Sergi Elizalde, Ashley M. Laughney, and Samuel F. Bakhoum. A Markov chain for numerical chromosomal instability in clonally expanding populations. PLoS computational biology, 14(9):e1006447, 2018.

[35] Ashley M. Laughney, Sergi Elizalde, Giulio Genovese, and Samuel F. Bakhoum. Dynamics of Tumor Heterogeneity Derived from Clonal Karyotypic Evolution. Cell Reports, 12(5):809–820, August 2015.

[36] Richard S Varga. Geršgorin and his circles, volume 36. Springer Science & Business Media, 2010.

[37] Dongqing Sun, Jin Wang, Ya Han, Xin Dong, Jun Ge, Rongbin Zheng, Xiaoying Shi, Binbin Wang, Ziyi Li, Pengfei Ren, Liangdong Sun, Yilv Yan, Peng Zhang, Fan Zhang, Taiwen Li, and Chenfei Wang. TISCH: a comprehensive web resource enabling interactive single-cell transcriptome visualization of tumor microenvironment. Nucleic Acids Research, 49(D1):D1420–D1430, January 2021.

[38] Charles P. Couturier, Shamini Ayyadhury, Phuong U. Le, Javad Nadaf, Jean Monlong, Gabriele Riva, Redouane Allache, Salma Baig, Xiaohua Yan, Mathieu Bourgey, Changseok Lee, Yu Chang David Wang, V. Wee Yong, Marie-Christine Guiot, Hamed Najafabadi, Bratislav Misic, Jack Antel, Guillaume Bourque, Jiannis Ragoussis, and Kevin Petrecca. Single-cell RNA-seq reveals that glioblastoma recapitulates a normal neurodevelopmental hierarchy. Nature Communications, 11(1):3406, July 2020. Number: 1 Publisher: Nature Publishing Group.

[39] Ruixi Li, Bo Liao, Bo Wang, Chan Dai, Xin Liang, Geng Tian, and Fangxiang Wu. Identification of Tumor Tissue of Origin with RNA-Seq Data and Using Gradient Boosting Strategy. BioMed Research International, 2021:6653793, February 2021.

[40] Niha Beig, Jay Patel, Prateek Prasanna, Virginia Hill, Amit Gupta, Ramon Correa, Kaustav Bera, Salendra Singh, Sasan Partovi, Vinay Varadan, Manmeet Ahluwalia, Anant Madabhushi, and Pallavi Tiwari. Radiogenomic analysis of hypoxia pathway is predictive of overall survival in Glioblastoma. Scientific Reports, 8(1):7, January 2018.

[41] Qinghua Xu, Jinying Chen, Shujuan Ni, Cong Tan, Midie Xu, Lei Dong, Lin Yuan, Qifeng Wang, and Xiang Du. Pan-cancer transcriptome analysis reveals a gene expression signature for the identification of tumor tissue origin. Modern Pathology, 29(6):546–556, June 2016. Number: 6 Publisher: Nature Publishing Group.

[42] Zhen Yang, Andrew Wong, Diana Kuh, Dirk S. Paul, Vardhman K. Rakyan, R. David Leslie, Shijie C. Zheng, Martin Widschwendter, Stephan Beck, and Andrew E. Teschendorff. Correlation of an epigenetic mitotic clock with cancer risk. Genome Biology, 17:205, October 2016.

[43] Noemi Andor, Billy T. Lau, Claudia Catalanotti, Anuja Sathe, Matthew A. Kubit, Jiamin Chen, Susan M. Grimes, Carlos Jose Suarez, and Hanlee P. Ji. Joint single cell DNA-seq and RNA-seq of gastric cancer cell lines reveals rules of in vitro evolution. NAR Genomics and Bioinformatics (in press), March 2020.

[44] Roel G W Verhaak, Katherine A Hoadley, Elizabeth Purdom, Victoria Wang, Yuan Qi, Matthew D Wilkerson, C Ryan Miller, Li Ding, Todd Golub, Jill P Mesirov, Gabriele Alexe, Michael Lawrence, Michael O’Kelly, Pablo Tamayo, Barbara A Weir, Stacey Gabriel, Wendy Winckler, Supriya Gupta, Lakshmi Jakkula, Heidi S Feiler, J Graeme Hodgson, C David James, Jann N Sarkaria, Cameron Brennan, Ari Kahn, Paul T Spellman, Richard K Wilson, Terence P Speed, Joe W Gray, Matthew Meyerson, Gad Getz, Charles M Perou, D Neil Hayes, and Cancer Genome Atlas Research Network. Integrated genomic analysis identifies clinically relevant subtypes of glioblastoma characterized by abnormalities in PDGFRA, IDH1, EGFR, and NF1. Cancer cell, 17(1):98–110, January 2010.

[45] Uri Ben-David, Benjamin Siranosian, Gavin Ha, Helen Tang, Yaara Oren, Kunihiko Hinohara, Craig A. Strathdee, Joshua Dempster, Nicholas J. Lyons, Robert Burns, Anwesha Nag, Guillaume Kugener, Beth Cimini, Peter Tsvetkov, Yosef E. Maruvka, Ryan O’Rourke, Anthony Garrity, Andrew A. Tubelli, Pratiti Bandopadhayay, Aviad Tsherniak, Francisca Vazquez, Bang Wong, Chet Birger, Mahmoud Ghandi, Aaron R. Thorner, Joshua A. Bittker, Matthew Meyerson, Gad Getz, Rameen Beroukhim, and Todd R. Golub. Genetic and transcriptional evolution alters cancer cell line drug response. Nature, 560(7718):325–330, August 2018.

[46] Uri Ben-David, Gavin Ha, Yuen-Yi Tseng, Noah F. Greenwald, Coyin Oh, Juliann Shih, James M. McFarland, Bang Wong, Jesse S. Boehm, Rameen Beroukhim, and Todd R. Golub. Patient-derived xenografts undergo mouse-specific tumor evolution. Nature Genetics, 49(11):1567–1575, November 2017.

[47] Samuel F. Bakhoum, Bryan Ngo, Ashley M. Laughney, Julie-Ann Cavallo, Charles J. Murphy, Peter Ly, Pragya Shah, Roshan K. Sriram, Thomas B. K. Watkins, Neil K. Taunk, Mercedes Duran, Chantal Pauli, Christine Shaw, Kalyani Chadalavada, Vinagolu K. Rajasekhar, Giulio Genovese, Subramanian Venkatesan, Nicolai J. Birkbak, Nicholas McGranahan, Mark Lundquist, Quincey LaPlant, John H. Healey, Olivier Elemento, Christine H. Chung, Nancy Y. Lee, Marcin Imielenski, Gouri Nanjangud, Dana Pe’er, Don W. Cleveland, Simon N. Powell, Jan Lammerding, Charles Swanton, and Lewis C. Cantley. Chromosomal instability drives metastasis through a cytosolic DNA response. Nature, 553(7689):467–472, January 2018.

[48] Sonja Hänzelmann, Robert Castelo, and Justin Guinney. GSVA: gene set variation analysis for microarray and RNA-Seq data. BMC Bioinformatics, 14(1):7, January 2013.

[49] David Croft, Antonio Fabregat Mundo, Robin Haw, Marija Milacic, Joel Weiser, Guanming Wu, Michael Caudy, Phani Garapati, Marc Gillespie, Maulik R. Kamdar, Bijay Jassal, Steven Jupe, Lisa Matthews, Bruce May, Stanislav Palatnik, Karen Rothfels, Veronica Shamovsky, Heeyeon Song, Mark Williams, Ewan Birney, Henning Hermjakob, Lincoln Stein, and Peter D’Eustachio. The Reactome pathway knowledgebase. Nucleic Acids Research, 42(Database issue):D472–D477, January 2014.

[50] DA Rew and GD Wilson. Cell production rates in human tissues and tumours and their significance. Part II: clinical data. European Journal of Surgical Oncology (EJSO), 26(4):405–417, 2000. Publisher: Elsevier.

[51] Karin Haustermans, Lucien Vanuytsel, Karel Geboes, T Lerut, J Van Thillo, J Leysen, Willy Coosemans, and E Van Der Schueren. In vivo cell kinetic measurements in human oesophageal cancer: what can be learned from multiple biopsies? European Journal of Cancer, 30(12):1787–1791, 1994. Publisher: Elsevier.

[52] Seung Joon Choi, Hyung-Sik Kim, Su-Joa Ahn, Yu Mi Jeong, and Hye-Young Choi. Evaluation of the growth pattern of carcinoma of colon and rectum by MDCT. Acta Radiologica, 54(5):487–492, 2013. Publisher: SAGE Publications Sage UK: London, England.

[53] Mitsunobu IDE, Minoru Jimbo, Masaaki Yamamoto, Yutaka Umebara, Shinji Hagiwara, and Osami Kubo. Growth rate of intracranial meningioma: tumor doubling time and proliferating cell nuclear antigen staining index. Neurologia medico-chirurgica, 35(5):289–293, 1995. Publisher: The Japan Neurosurgical Society.

[54] S Bolin, E Nilsson, and R Sjödahl. Carcinoma of the colon and rectum–growth rate. Annals of surgery, 198(2):151, 1983. Publisher: Lippincott, Williams, and Wilkins.

[55] RG Margolese and Buchholz TA HG. Natural history and prognostic markers. Holland-Frei Cancer Medicine. 6th edition ed. Hamilton (ON): BC Decker, 2003.

[56] GM Zharinov and VA Gushchin. The rate of tumor growth and cell loss in cervical cancer. Voprosy onkologii, 35(1):21–25, 1989.

[57] J Andrew Carlson. Tumor doubling time of cutaneous melanoma and its metastasis. The American journal of dermatopathology, 25(4):291–299, 2003. Publisher: LWW.

[58] Kassem Harris, Inga Khachaturova, Basem Azab, Theodore Maniatis, Srujitha Murukutla, Michel Chalhoub, Hassan Hatoum, Thomas Kilkenny, Dany Elsayegh, Rabih Maroun, and others. Small cell lung cancer doubling time and its effect on clinical presentation: a concise review. Clinical Medicine Insights: Oncology, 6:CMO–S9633, 2012. Publisher: SAGE Publications Sage UK: London, England.

[59] Katja Roesch, Dirk Hasenclever, and Markus Scholz. Modelling lymphoma therapy and outcome. Bulletin of mathematical biology, 76(2):401–430, 2014. Publisher: Springer.

[60] Lijun Sun, Jiaxi Wu, Fenghe Du, Xiang Chen, and Zhijian J. Chen. Cyclic GMP-AMP synthase is a cytosolic DNA sensor that activates the type I interferon pathway. Science (New York, N.Y.), 339(6121):786–791, February 2013.

[61] Yuk Yuen Lan, Diana Londoño, Richard Bouley, Michael S. Rooney, and Nir Hacohen. Dnase2a deficiency uncovers lysosomal clearance of damaged nuclear DNA via autophagy. Cell Reports, 9(1):180–192, October 2014.

[62] Karen J. Mackenzie, Paula Carroll, Carol-Anne Martin, Olga Murina, Adeline Fluteau, Daniel J. Simpson, Nelly Olova, Hannah Sutcliffe, Jacqueline K. Rainger, Andrea Leitch, Ruby T. Osborn, Ann P. Wheeler, Marcin Nowotny, Nick Gilbert, Tamir Chandra, Martin A. M. Reijns, and Andrew P. Jackson. cGAS surveillance of micronuclei links genome instability to innate immunity. Nature, 548(7668):461–465, August 2017.

[63] Uri Ben-David and Angelika Amon. Context is everything: aneuploidy in cancer. Nature Reviews. Genetics, 21(1):44–62, January 2020.

[64] Kristian Cibulskis, Michael S. Lawrence, Scott L. Carter, Andrey Sivachenko, David Jaffe, Carrie Sougnez, Stacey Gabriel, Matthew Meyerson, Eric S. Lander, and Gad Getz. Sensitive detection of somatic point mutations in impure and heterogeneous cancer samples. Nature Biotechnology, 31(3):213–219, 2013. 00000.

[65] Zuzana Storchova and Christian Kuffer. The consequences of tetraploidy and aneuploidy. Journal of Cell Science, 121(23):3859–3866, December 2008. Publisher: The Company of Biologists Ltd Section: Commentary.

[66] Ankit Shukla, Thu H. M. Nguyen, Sarat B. Moka, Jonathan J. Ellis, John P. Grady, Harald Oey, Alexandre S. Cristino, Kum Kum Khanna, Dirk P. Kroese, Lutz Krause, Eloise Dray, J. Lynn Fink, and Pascal H. G. Duijf. Chromosome arm aneuploidies shape tumour evolution and drug response. Nature Communications, 11(1):449, January 2020. Number: 1 Publisher: Nature Publishing Group.

[67] Salehi S, Kabeer F, Ceglia N, Andronescu M, Williams Mj, Campbell Kr, Masud T, Wang B, Biele J, Brimhall J, Gee D, Lee H, Ting J, Zhang Aw, Tran H, O’Flanagan C, Dorri F, Rusk N, de Algara Tr, Lee Sr, Cheng Byc, Eirew P, Kono T, Pham J, Grewal D, Lai D, Moore R, Mungall Aj, Marra Ma, undefined, McPherson A, Bouchard-Côté A, Aparicio S, and Shah Sp. Clonal fitness inferred from time-series modelling of single-cell cancer genomes. Nature, 595(7868):585–590, June 2021.

[68] Bastien Nguyen, Christopher Fong, Anisha Luthra, Shaleigh A. Smith, Renzo G. DiNatale, Subhiksha Nandakumar, Henry Walch, Walid K. Chatila, Ramyasree Madupuri, Ritika Kundra, Craig M. Bielski, Brooke Mastrogiacomo, Mark T. A. Donoghue, Adrienne Boire, Sarat Chandarlapaty, Karuna Ganesh, James J. Harding, Christine A. Iacobuzio-Donahue, Pedram Razavi, Ed Reznik, Charles M. Rudin, Dmitriy Zamarin, Wassim Abida, Ghassan K. Abou-Alfa, Carol Aghajanian, Andrea Cercek, Ping Chi, Darren Feldman, Alan L. Ho, Gopakumar Iyer, Yelena Y. Janjigian, Michael Morris, Robert J. Motzer, Eileen M. O’Reilly, Michael A. Postow, Nitya P. Raj, Gregory J. Riely, Mark E. Robson, Jonathan E. Rosenberg, Anton Safonov, Alexander N. Shoushtari, William Tap, Min Yuen Teo, Anna M. Varghese, Martin Voss, Rona Yaeger, Marjorie G. Zauderer, Nadeem Abu-Rustum, Julio Garcia-Aguilar, Bernard Bochner, Abraham Hakimi, William R. Jarnagin, David R. Jones, Daniela Molena, Luc Morris, Eric Rios-Doria, Paul Russo, Samuel Singer, Vivian E. Strong, Debyani Chakravarty, Lora H. Ellenson, Anuradha Gopalan, Jorge S. Reis-Filho, Britta Weigelt, Marc Ladanyi, Mithat Gonen, Sohrab P. Shah, Joan Massague, Jianjiong Gao, Ahmet Zehir, Michael F. Berger, David B. Solit, Samuel F. Bakhoum, Francisco Sanchez-Vega, and Nikolaus Schultz. Genomic characterization of metastatic patterns from prospective clinical sequencing of 25,000 patients. Cell, 185(3):563–575.e11, February 2022.

[69] Christopher Abbosh, Nicolai J. Birkbak, Gareth A. Wilson, Mariam Jamal-Hanjani, Tudor Constantin, Raheleh Salari, John Le Quesne, David A. Moore, Selvaraju Veeriah, Rachel Rosenthal, Teresa Marafioti, Eser Kirkizlar, Thomas B. K. Watkins, Nicholas McGranahan, Sophia Ward, Luke Martinson, Joan Riley, Francesco Fraioli, Maise Al Bakir, Eva Grönroos, Francisco Zambrana, Raymondo Endozo, Wenya Linda Bi, Fiona M. Fennessy, Nicole Sponer, Diana Johnson, Joanne Laycock, Seema Shafi, Justyna Czyzewska-Khan, Andrew Rowan, Tim Chambers, Nik Matthews, Samra Turajlic, Crispin Hiley, Siow Ming Lee, Martin D. Forster, Tanya Ahmad, Mary Falzon, Elaine Borg, David Lawrence, Martin Hayward, Shyam Kolvekar, Nikolaos Panagiotopoulos, Sam M. Janes, Ricky Thakrar, Asia Ahmed, Fiona Blackhall, Yvonne Summers, Dina Hafez, Ashwini Naik, Apratim Ganguly, Stephanie Kareht, Rajesh Shah, Leena Joseph, Anne Marie Quinn, Phil A. Crosbie, Babu Naidu, Gary Middleton, Gerald Langman, Simon Trotter, Marianne Nicolson, Hardy Remmen, Keith Kerr, Mahendran Chetty, Lesley Gomersall, Dean A. Fennell, Apostolos Nakas, Sridhar Rathinam, Girija Anand, Sajid Khan, Peter Russell, Veni Ezhil, Babikir Ismail, Melanie Irvin-Sellers, Vineet Prakash, Jason F. Lester, Malgorzata Kornaszewska, Richard Attanoos, Haydn Adams, Helen Davies, Dahmane Oukrif, Ayse U. Akarca, John A. Hartley, Helen L. Lowe, Sara Lock, Natasha Iles, Harriet Bell, Yenting Ngai, Greg Elgar, Zoltan Szallasi, Roland F. Schwarz, Javier Herrero, Aengus Stewart, Sergio A. Quezada, Karl S. Peggs, Peter Van Loo, Caroline Dive, C. Jimmy Lin, Matthew Rabinowitz, Hugo J. W. L. Aerts, Allan Hackshaw, Jacqui A. Shaw, Bernhard G. Zimmermann, TRACERx consortium, PEACE consortium, and Charles Swanton. Phylogenetic ctDNA analysis depicts early-stage lung cancer evolution. Nature, 545(7655):446–451, April 2017.

[70] Mariam Jamal-Hanjani, Gareth A. Wilson, Nicholas McGranahan, Nicolai J. Birkbak, Thomas B. K. Watkins, Selvaraju Veeriah, Seema Shafi, Diana H. Johnson, Richard Mitter, Rachel Rosenthal, Max Salm, Stuart Horswell, Mickael Escudero, Nik Matthews, Andrew Rowan, Tim Chambers, David A. Moore, Samra Turajlic, Hang Xu, Siow-Ming Lee, Martin D. Forster, Tanya Ahmad, Crispin T. Hiley, Christopher Abbosh, Mary Falzon, Elaine Borg, Teresa Marafioti, David Lawrence, Martin Hayward, Shyam Kolvekar, Nikolaos Panagiotopoulos, Sam M. Janes, Ricky Thakrar, Asia Ahmed, Fiona Blackhall, Yvonne Summers, Rajesh Shah, Leena Joseph, Anne M. Quinn, Phil A. Crosbie, Babu Naidu, Gary Middleton, Gerald Langman, Simon Trotter, Marianne Nicolson, Hardy Remmen, Keith Kerr, Mahendran Chetty, Lesley Gomersall, Dean A. Fennell, Apostolos Nakas, Sridhar Rathinam, Girija Anand, Sajid Khan, Peter Russell, Veni Ezhil, Babikir Ismail, Melanie Irvin-Sellers, Vineet Prakash, Jason F. Lester, Malgorzata Kornaszewska, Richard Attanoos, Haydn Adams, Helen Davies, Stefan Dentro, Philippe Taniere, Brendan O’Sullivan, Helen L. Lowe, John A. Hartley, Natasha Iles, Harriet Bell, Yenting Ngai, Jacqui A. Shaw, Javier Herrero, Zoltan Szallasi, Roland F. Schwarz, Aengus Stewart, Sergio A. Quezada, John Le Quesne, Peter Van Loo, Caroline Dive, Allan Hackshaw, Charles Swanton, and TRACERx Consortium. Tracking the Evolution of Non-Small-Cell Lung Cancer. The New England Journal of Medicine, 376(22):2109–2121, June 2017.

[71] Ruli Gao, Shanshan Bai, Ying C. Henderson, Yiyun Lin, Aislyn Schalck, Yun Yan, Tapsi Kumar, Min Hu, Emi Sei, Alexander Davis, Fang Wang, Simona F. Shaitelman, Jennifer Rui Wang, Ken Chen, Stacy Moulder, Stephen Y. Lai, and Nicholas E. Navin. Delineating copy number and clonal substructure in human tumors from single-cell transcriptomes. Nature Biotechnology, 39(5):599–608, May 2021. Number: 5 Publisher: Nature Publishing Group.

[72] Jiqiu Cheng, Jonas Demeulemeester, David C. Wedge, Hans Kristian M. Vollan, Jason J. Pitt, Hege G. Russnes, Bina P. Pandey, Gro Nilsen, Silje Nord, Graham R. Bignell, Kevin P. White, Anne-Lise Børresen-Dale, Peter J. Campbell, Vessela N. Kristensen, Michael R. Stratton, Ole Christian Lingjærde, Yves Moreau, and Peter Van Loo. Pan-cancer analysis of homozygous deletions in primary tumours uncovers rare tumour suppressors. Nature Communications, 8(1):1221, October 2017.

[73] Paul Little, Dan-Yu Lin, and Wei Sun. Associating somatic mutations to clinical outcomes: a pan-cancer study of survival time. Genome Medicine, 11(1):37, May 2019.

[74] Andor, Noemi, Altrock, Philipp, Jain, Navami, and Gomes, Ana. Tipping cancer cells over the edge: the context-dependent cost of high ploidy. Cancer Research. In press., November 2021.

[75] Ron Sender and Ron Milo. The distribution of cellular turnover in the human body. Nature Medicine, 27(1):45–48, January 2021. Number: 1 Publisher: Nature Publishing Group.

[76] Neil J Ganem, Zuzana Storchova, and David Pellman. Tetraploidy, aneuploidy and cancer. Current opinion in genetics & development, 17(2):157–162, 2007. Publisher: Elsevier.

[77] Karen Crasta, Neil J Ganem, Regina Dagher, Alexandra B Lantermann, Elena V Ivanova, Yunfeng Pan, Luigi Nezi, Alexei Protopopov, Dipanjan Chowdhury, and David Pellman. DNA breaks and chromosome pulverization from errors in mitosis. Nature, 482(7383):53–58, 2012. Publisher: Nature Publishing Group.

[78] Cheng-Zhong Zhang, Alexander Spektor, Hauke Cornils, Joshua M Francis, Emily K Jackson, Shiwei Liu, Matthew Meyerson, and David Pellman. Chromothripsis from DNA damage in micronuclei. Nature, 522(7555):179–184, 2015. Publisher: Nature Publishing Group.

[79] Emily M Hatch, Andrew H Fischer, Thomas J Deerinck, and Martin W Hetzer. Catastrophic nuclear envelope collapse in cancer cell micronuclei. Cell, 154(1):47–60, 2013. Publisher: Elsevier.

[80] Jake C Forster, Michael JJ Douglass, Wendy M Harriss-Phillips, and Eva Bezak. Simulation of head and neck cancer oxygenation and doubling time in a 4D cellular model with angiogenesis. Scientific reports, 7(1):1–11, 2017. Publisher: Nature Publishing Group.

[81] Ryan J. Quinton, Amanda DiDomizio, Marc A. Vittoria, Kristýna Kotýnková, Carlos J. Ticas, Sheena Patel, Yusuke Koga, Jasmine Vakhshoorzadeh, Nicole Hermance, Taruho S. Kuroda, Neha Parulekar, Alison M. Taylor, Amity L. Manning, Joshua D. Campbell, Neil J. Ganem. Whole genome doubling confers unique genetic vulnerabilities on tumor cells. Nature, 590(7846):492–497, February 2021.

