## Supplementary Figures and Tables for "Intra-tumor heterogeneity, turnover rate and karyotype space shape susceptibility to missegregation-induced extinction"

### Supplementary Information

#### 1.1 Numerical methods for calculating the critical curve

Critical curves are derived by leveraging the fact that the critical turnover rate for a given $\beta$ is when the rate of missegregation induced cell death equals the growth rate $(\lambda - \mu)$. Let $J$ be the Jacobian defined in (9). The system reaches a steady state when $n_i J = 0$. Nontrivial solutions can be found by choosing functions for the mis-segregation rate $\beta$ and death rate $\mu$ (which parameterise the matrix $J$) such that the dominant eigenvalue is zero. A robust method for determining pairs of values $(B, M)$ on the critical curve (which can cope with heterogeneous functions for $\beta, \mu$), is to: 1) choose an input value $B$ for the function $\beta(i, B)$, then 2) find the value of $M$ parameterizing $\mu(i, M)$ such that the dominant eigenvalue of J is equal to zero. The eigenvector corresponding to this eigenvalue is the steady state of the system. Although this method is robust, it requires multiple generations of the matrix $J$ with different values of $M$ and so can become slow when $J$ is a large matrix. A considerable time saving can be achieved when death rate does not depend on copy number state, i.e. $\mu = M$. In that case, $J$ can be expressed in terms of the adjusted matrix $J^*$:

$$J = J^* + \mu I, \tag{15}$$

where $J^*$ represents the transition matrix with $\mu = 0$ and $I$ is the identity matrix. It can be shown that the eigenvectors of $J$ are identical to those of $J^*$, therefore the steady state population distribution (i.e. the karyotype composition) does not depend on the death rate $\mu$ and is given by the dominant eigenvector of $J^*$. Denoting this eigenvector $\bar{n}_i$, after normalising $\bar{n}_i$ such that the entries sum to one, $\sum_i \bar{n}_i J^*$ gives the net growth rate of the cell population when $\mu = 0$ and the population *distribution* has reached steady state (although net population growth still occurs when $\mu = 0$). Therefore, the true steady state (and therefore the critical curve) is found by choosing the value of $\mu$ which exactly negates this growth, i.e: $\mu = \sum_i \bar{n}_i J^*$. We note that if death rates are heterogeneous then $J^*$ is no longer independent of the death rate $\mu$, i.e. the approach relies on the assumption of homogeneous rates.

Both of the above approaches work when a single chromosome type has a finite viable copy number interval as well as when multiple chromosomes have finite intervals (i.e. $n_i$ becomes $n_{\vec{i}}$). However both become infeasible when we have $k \gg 1$ chromosome types, each with multiple copies, as the matrix $J$ becomes too large. However the critical curve can still be generated by extrapolating from the single chromosome case. Summing over the rows of the matrix $q_{ij}$ (eq. (6)) gives a probability vector $q_i^*$, which contains the probability that a daughter of a cell with $i$ chromosome copies receives a viable number of copies. The overall probability that a daughter cell from the steady state population receives an unviable number of copies for a single chromosome type is therefore: $q_d = 1 - q_i^* \cdot \bar{n}_i$. The probability that a daughter receives viable copy numbers for all $k$ chromosome types is: $q_a^k = (1 - q_d)^k = (q_i^* \cdot \bar{n}_i)^k$. The rate of change of the alive cell population is given by:

$$\frac{dn_a}{dt} = \lambda n_a (2q_a^k - 1) - \mu n_a, \tag{16}$$

,

where $n_a$ is the total number of alive cells. Setting to zero, we have $\mu = 2q_a - 1$.

##### 1.2 MIE is impossible with the number of viable karyotypes $K \to \infty$.

Let us assume that no more than one copy of a critical chromosome can mis-segregate during one cell division, then $\mathbf{q}$ has elements

$$q_{ij} = \begin{cases} 1-\beta & \text{if } i = j, \\ \beta & \text{if } |i-j| = 1, \\ 0 & \text{otherwise}, \end{cases} \tag{17}$$

If all parameters are independent of karyotype and with the number of compartments $K \to \infty$, our Jacobian becomes:

$$J_{ij} = \begin{cases} \lambda_i(1-2\beta_i) - \mu_i & \text{if } i = j, \\ \lambda_i\beta_i & \text{if } i = j+1, \\ \lambda_j\beta_j & \text{if } i = j-1, \\ 0 & \text{otherwise}. \end{cases}$$

This has the structure of a Toeplitz matrix, for which there has been a lot of theory developed over the past 100 years. We first assume the simple case that all parameters are equal, then defining $B = J - \psi I$ where $\psi$ is the eigenvalues of $J$, we can define $\phi = \lambda(1-2\beta) - \mu - \psi$ and $\alpha = \lambda\beta$. Thus

$$B_{ij} = \begin{cases} \phi & \text{if } i = j, \\ \alpha & \text{if } |i-j| = 1, \\ 0 & \text{otherwise}. \end{cases}$$

Define the characteristic polynomial of $B_K$ to be $F_K(\phi)$ where $K$ is for the dimension of the matrix. The roots of $F_K(\phi) = 0$ will be the eigenvalues. Using the Laplace expansion for computing the determinant, we can set up a recursion relation for $F_K$:

$$F_K = \phi F_{K-1} - \alpha^2 F_{K-2}, \quad F_0 = 1, \; F_{-1} = 0. \tag{18}$$

This is a linear difference equation and can be solved with the Ansatz $F_K = z^k$. We obtain

$$z_{\pm} = \frac{\phi}{2}\left(1 \pm \sqrt{1-\beta^2}\right),$$

where $\beta = 2\alpha/\phi$. Suppose first that $\beta^2 < 1$, then we have

$$F_K = c_1 z_+^K + c_2 z_-^K.$$

The initial conditions imply

$$F_K = \frac{z_+^{K+1} - z_-^{K+1}}{z_+ - z_-}.$$

Now, recall our condition $\beta^2 < 1$, which implies that $2\alpha < |\phi|$. If the eigenvalue is large, this number is negative so this relation becomes $\lambda - \mu < \psi$. Since we assume that $\lambda > \mu$, we see that $\psi > 0$ and so the largest eigenvalue will certainly be positive and so we will not have MIE.

Instead, suppose that $\beta^2 > 1$, then we can rewrite the characteristic polynomial as

$$F_K = \frac{\alpha^K \sin\left[(K+1)\arccos\left(\frac{\phi}{2\alpha}\right)\right]}{\sin\left[\arccos\left(\frac{\phi}{2\alpha}\right)\right]}.$$

Setting this to 0 implies that

$$\phi_m = 2\alpha\cos\left(\frac{m\pi}{K+1}\right),\ m = 1, \ldots, K$$

are the solutions for $\phi$. Thus the eigenvalues can be found exactly in this case and are given by

$$\psi_m = \lambda(1-2\beta) - \mu - \phi_m. \tag{19}$$

The maximum eigenvalue is when $\phi_m$ is the smallest which occurs at $m = K$. In the limit $K \to \infty$

$$\cos\left(\frac{K\pi}{K+1}\right) = -\cos\left(\frac{\pi}{K+1}\right) = -\left[1 - \frac{\pi^2}{2(K+1)^2} + \mathcal{O}(K^{-4})\right].$$

Thus the critical curve (when the maximum eigenvalue is 0) in the limit $K \to \infty$ is given by

$$\beta_c = \frac{2(K+1)^2}{\pi^2}\left(1 - \frac{\mu}{\lambda}\right). \tag{20}$$

Recall that $\beta \in [0,1]$ and so for sufficiently large $K$, this will push us outside of this bound. Thus, we have shown that if all parameters are equal, and jumps of size one are permitted, then as $K \to \infty$, MIE is not possible. The above result is rigorous. To discuss the general case, a heuristic argument in the continuum limit shows that MIE is impossible if the set of viable regions tends to infinity (Supplementary Information 1.3). This also includes the possibility of regions of karyotype viability interspersed between nonviable regions.

#### 1.3 MIE is impossible for infinite compartments - continuum argument

For small jumps and variable coefficients, we appeal to the continuum model for the behavior. Expanding equation (8) for small jumps we have

$$\frac{dn_i}{dt} = \lambda_{i+1}\beta_{i+1}n_{i+1} - 2\lambda_i\beta_i n_i + \lambda_{i-1}\beta_{i-1}n_{i-1} + (\lambda_i - \mu_i)n_i. \tag{21}$$

Define $p = i/K$, and $\Delta p = 1/K$. For $K \to \infty$ we have

$$\lambda_{i+1}\beta_{i+1}n_{i+1} - 2\lambda_i\beta_i n_i + \lambda_{i-1}\beta_{i-1}n_{i-1} \to \frac{1}{K^2}\frac{\partial^2}{\partial p^2}(\beta\lambda n). \tag{22}$$

This leads to

$$\frac{\partial n}{\partial t} = (\lambda - \mu)n + \frac{1}{K^2}\frac{\partial^2}{\partial p^2}(\beta\lambda n), \tag{23}$$

where $p \in (0,1)$ and $n(0) = n(1) = 0$. We make use of the Ansatz, $n = e^{\alpha t}U(p)$, which results in the eigenvalue problem

$$\alpha U = \frac{1}{K^2}(\beta\lambda U)'' + (\lambda - \mu)U. \tag{24}$$

For simplicity, we define $q = \frac{1}{K^2}\beta\lambda$ and convert to Sturm-Liouville form

$$\begin{aligned} \alpha U &= qU'' + 2q'U' + (\lambda - \mu + q'')U \\ \alpha q U &= (q^2U')' + q(\lambda - \mu + q'')U. \end{aligned}$$

Multiplying by $U$ and integrating over $p$, we obtain an equation for $\alpha$ (the Rayleigh Quotient):

$$\alpha = \frac{\int_0^1 -q^2(U')^2 + q(\lambda - \mu + q'')U^2 \; dp}{\int_0^1 qU^2 \; dp} = \frac{\int_0^1 -\frac{(\beta\lambda)^2}{K^2}(U')^2 + \beta\lambda\left[\lambda - \mu + \frac{(\beta\lambda)''}{K^2}\right]U^2 \; dp}{\int_0^1 \beta\lambda U^2 \; dp}. \tag{25}$$

For $K \to \infty$ we may neglect the terms with $K^{-2}$ and obtain

$$\alpha = \frac{\int_0^1 \beta\lambda(\lambda - \mu)U^2 \; dp}{\int_0^1 \beta\lambda U^2 \; dp}. \tag{26}$$

Provided that $r = \lambda - \mu > 0$ for all $p$ we define $m = \inf(\beta\lambda(\lambda - \mu))$ and $P = \sup(\beta\lambda)$, it follows that

$$\alpha \geq \frac{m}{P} > 0.$$

Finally, we note that the Rayleigh quotient satisfies $\alpha_{\min} \leq \alpha \leq \alpha_{\max}$ for all amendable test functions $U$, where $\alpha_{\min}, \alpha_{\max}$ are the smallest and largest eigenvalues, respectively. Hence, $\alpha_{\max} > 0$ and so MIE is impossible.

Note that the proof did not rely on the whole integral having $r > 0$. Suppose that we have an almost everywhere differentiable function $U$ that is 0 on $[0,1]\backslash E$, with $E \subset [0,1]$ and $m, P$ defined as above. Then (26) becomes:

$$\alpha = \frac{\int_E \beta\lambda(\lambda - \mu)U^2 \; dp}{\int_E \beta\lambda U^2 \; dp} \geq \frac{m}{P} > 0, \tag{27}$$

provided that $|E| > 0$.

One final note, this may seem to imply that we have shown that a *finite* number of compartments in the small jump case can't have MIE, but this would be false. We must take a closer look at the passage to equilibrium. For simplicity suppose the viable karyotype interval is at the beginning $i = 1, \ldots, f(K)$ where $r > 0$. Defining $p = i/K$ as before, note that the viable interval length is on the order of $f(K)/K$. For the measure of this interval to remain finite, we require that $f(K) \to cK$ for $K \to \infty$ for some $0 < c < 1$ to rule out MIE. Essentially, the viable interval length must grow at the same order as the number of viable intervals grows, in order to be able to rule out MIE using this method.

#### 1.4 Ruling out MIE

Here, we establish sufficient conditions for MIE to not occur. These are based on Gershgorin's circle (GC) theorem which states that for matrix $A$ with elements $a_{ij}$, the eigenvalues $\psi$ satisfy:

$$|\psi - a_{ii}| \leq \sum_{j \neq i} |a_{ij}| = R_i. \tag{28}$$

Put differently, every eigenvalue of $A$ lies within at least one of the discs with center $a_{ii}$ and radius $R_i$.

Since MIE can be evaded if the maximum eigenvalue exceeds 0, a sufficient condition is that none of Gershgorin's circle contain a part of the negative reals. That is $a_{ii} - R_i > 0$ for all $i$.

For example in the case of small jumps (Supplementary Information 1.2) with homogeneous (i.e. constant) rates, we showed that MIE is impossible as the number of compartments tended to infinity. This was regardless of the choice of parameters. Here we see that using GC, a sufficient condition to avoid MIE is given by

$$\beta < \beta_c = \frac{1}{4}\left(1 - \frac{\mu}{\lambda}\right). \tag{29}$$

Of course, we know that for large $K$, *no* missegregation rate will lead to MIE; this just provides a sufficient condition.

If $\lambda, \beta$ and $\mu$ vary with ploidy $i$, we can still obtain a sufficient condition to avoid MIE given by:

$$\lambda_i(1 - 3\beta_i) - \mu_i - \lambda_{i+1}\beta_{i+1} > 0, \text{ for all } i. \tag{30}$$

If $\beta$ is homogeneous, then this reduces to

$$\beta < \beta_c = \frac{1}{3 + \lambda_{i+1}/\lambda_i}\left(1 - \frac{\mu_i}{\lambda_i}\right), \text{ for all } i. \tag{31}$$

The general problem (i.e. any arbitrary relation between $\lambda, \beta$ and $\mu$) can be handled analogously, but conclusions can only be made for specific forms of $\mathbf{q}$. We require both (10)-(11). As both of these need to be positive, we can find the minimum of these, which will provide sufficient condition to escape MIE.

Let us consider a more complicated example to show the power of this method. Suppose that $\lambda, \mu$ are homogeneous, but that $q_{ij} = q(i \to j)$ obeys the following relationship

$$q(i \to i \pm h) = \kappa_i \beta^h, \ h = 0, 1, \ldots, i. \tag{32}$$

This $\mathbf{q}$ can be thought of as one that favors small jumps but all jumps are permitted. Let us suppose that there is no limit on the number of compartments, then:

$$1 = \sum_{h=0}^{i} q(i \to i + h) = \kappa_i \sum_{h=0}^{i} \beta^h = \frac{\kappa_i(1 - \beta^{i+1})}{1 - \beta}.$$

This implies the relation for $\kappa_i$:

$$\kappa_i = \frac{1 - \beta}{1 - \beta^{i+1}}. \tag{33}$$

We now look at the sum in (10) (note the symmetry $q(i \to i+h) = q(i \to i-h)$),

$$2\kappa_i \sum_{h=1}^{i} \beta^h = \frac{2\beta(1-\beta^i)}{1-\beta^{i+1}} \tag{34}$$

The sum in (11) is trickier since in this case we are looking at the possibilities $q(i+h \to i)$ and it is clear that $q(i+h \to i) \neq q(i-h \to i)$ (because any given ploidy state $i$ is more likely to emerge from missegregations in a higher ploidy cell than in a lower ploidy cell). We look at the first part of the sum first

$$\sum_{h=1}^{\infty} q(i+h \to i) = \sum_{h=1}^{\infty} \kappa_{i+h}\beta^h = (1-\beta)\sum_{h=1}^{\infty} \frac{\beta^h}{1-\beta^{i+h+1}} = \beta + \mathcal{O}(\beta^{3+i}),$$

valid when $\beta \ll 1$. The sum from the other direction is nonzero only if $h \geq \lceil i/2 \rceil$:

$$\sum_{h=\lceil i/2 \rceil}^{i-1} q(i-h \to i) = \sum_{h=\lceil i/2 \rceil}^{i-1} \kappa_{i-h}\beta^h = (1-\beta)\sum_{h=\lceil i/2 \rceil}^{i-1} \frac{\beta^h}{1-\beta^{i-h+1}} = \beta^{\lceil i/2 \rceil} - \beta^i + \mathcal{O}(\beta^{i+1}).$$

Therefore, the sum term in (11) ($\sum_{j \neq k} \lambda_j q_{jk}$) becomes:

$$\beta + \beta^{\lceil i/2 \rceil} - \beta^i + \mathcal{O}(\beta^{1+i}). \tag{35}$$

The bigger sum sets the threshold and is maximal for large $i$. With this choice of $\mathbf{q}$, it is (10) which is the dominant condition. Thus, a sufficient condition to escape MIE is given by:

$$\beta < \beta_c = \frac{1}{4}\left(1 - \frac{\mu}{\lambda}\right). \tag{36}$$

The appearance of this relation potentially suggests a fundamental limit on a kernel of this type. Indeed, it is not hard to show that for general $\mathbf{q}$ with $q_{ii} = 1 - \beta_i$, (10) yields

$$\beta_i < \frac{1}{4}\left(1 - \frac{\mu_i}{\lambda_i}\right), \text{ for all } i.$$

Since both (10) and (11) must be met, the above condition is the upper bound on the $\beta_i$'s required to evade MIE.

#### 1.5 Extensions to model whole-genome doubling and micronuclei formation

Though WGDs are technically feasible in the model (through an $i$ parent giving birth to $2i$ and 0 ploidy offspring), this ignores the biological mechanisms for how WGD events may occur. There are two mechanisms for WGD [74, 75] – endoreplication (i.e. completely skipping mitosis) and cytokinesis failure which often generates binucleated whole genome doubled cells. Chromosomes in the two nuclei can then merge the subsequent mitosis after nuclear envelope breakdown. We can modify equation (8) to account for WGD by introducing terms

$$\frac{dn_i}{dt} = \sum_j \lambda_j n_j q_{ji} - \lambda_i n_i (1 - q_{ii}) - \mu_i n_i - \theta_i n_i + \theta_{i/2} n_{i/2},$$

where $\theta_{i/2} = 0$ if $i/2$ is not an integer.

Mounting evidence suggests that a cell which mis-segregates many chromosomes has a higher risk to die than a cell mis-segregating only one chromosome [27, 28]. Our current model structure however does not distinguish between these scenarios. To account for this intolerance, we can amend the inflow term by introducing a karyotype transition coefficient $\phi_{ij}$. One possible form $\phi = (1+|i-j|)^\alpha$ with $\alpha < 0$, where only relative distance is important. Other forms that take the parental state into account could also be considered. Combining this with the above, leads to

$$\frac{dn_i}{dt} = \sum_j \phi_{ji} \lambda_j n_j q_{ji} - \lambda_i n_i (1 - q_{ii}) - \mu_i n_i - \theta_i n_i + \theta_{i/2} n_{i/2}.$$

The only requirement is that $\phi_{ij}$ must fall off with increasing distance between states.

Yet another assumption, which could be eventually relaxed is the ploidy conservation in equation (3). The errors during mitosis could lead to the formation of micronuclei which are not directly involved in further downwind mitosis [18, 36, 76–78]. This amounts to replacing equation (3) with $2i_k \geq j_k^{(1)} + j_k^{(2)}$ for all $k$.

Supplementary Table 1: ***In vivo* tumor dynamics across nine cancer types**. Birth rates taken from [49] and growth rates taken from multiple sources (cited in third column). The death rate is inferred. Many of the birth rates are comparable, but the net growth rates vary over orders of magnitude, which implies very different death rates. Last column shows percent contribution of respective normal human cell types to a total daily mass turnover of 80 g (taken from [79]) – these are correlated to both birth and death rates reported in tumors (Pearson $r \geq 0.923; P < 0.026$).

| Cancer type | birth rate ($\lambda$) [49] | growth rate ($r$) | death rate ($\mu = \lambda - r$) | normal turnover (%) |
|---|---|---|---|---|
| Head/Neck | 0.192 (0.015 - 1.667) | 0.007 (0.003 - 0.033) [80] | 0.185 (0 - 1.664) | NA |
| Esophageal | 0.207 (0.018 - 0.625) | 0.154 (0.035 - 0.347) [50] | 0.053 (0 - 0.59) | NA |
| Colorectal | 0.239 (0.019 - 1.111) | 0.003 (0.0003 - 0.038) [51] | 0.236 (0 - 1.111) | 41 [79] |
| Rectal | 0.303 (0.179 - 0.417) | 0.005 (0.0004 - 0.013) [53] | 0.298 (0.166 - 0.416) | 41 [79] |
| Breast | 0.122 (0.021 - 0.556) | 0.003 (0.0003 - 0.016) [54] | 0.119 (0.005 - 0.555) | NA |
| Cervix | 0.25 (0.159 - 0.323) | 0.002 (0.0004 - 0.012) [55] | 0.248 (0.147 - 0.322) | NA |
| Melanoma | 0.139 (0.024 - 0.286) | 0.007 (0.002 - 0.014) [56] | 0.132 (0.01 - 0.284) | 4 [79] |
| SC Lung | 0.122 (0.036 - 0.667) | 0.008 (0.005 - 0.013) [57] | 0.114 (0.023 - 0.662) | 0.5 [79] |
| D-LBCL | 0.063 (0.043 - 0.4) | 0.023 (0.01 - 0.5) [58] | 0.04 (0 - 0.39) | 1.7 [79] |

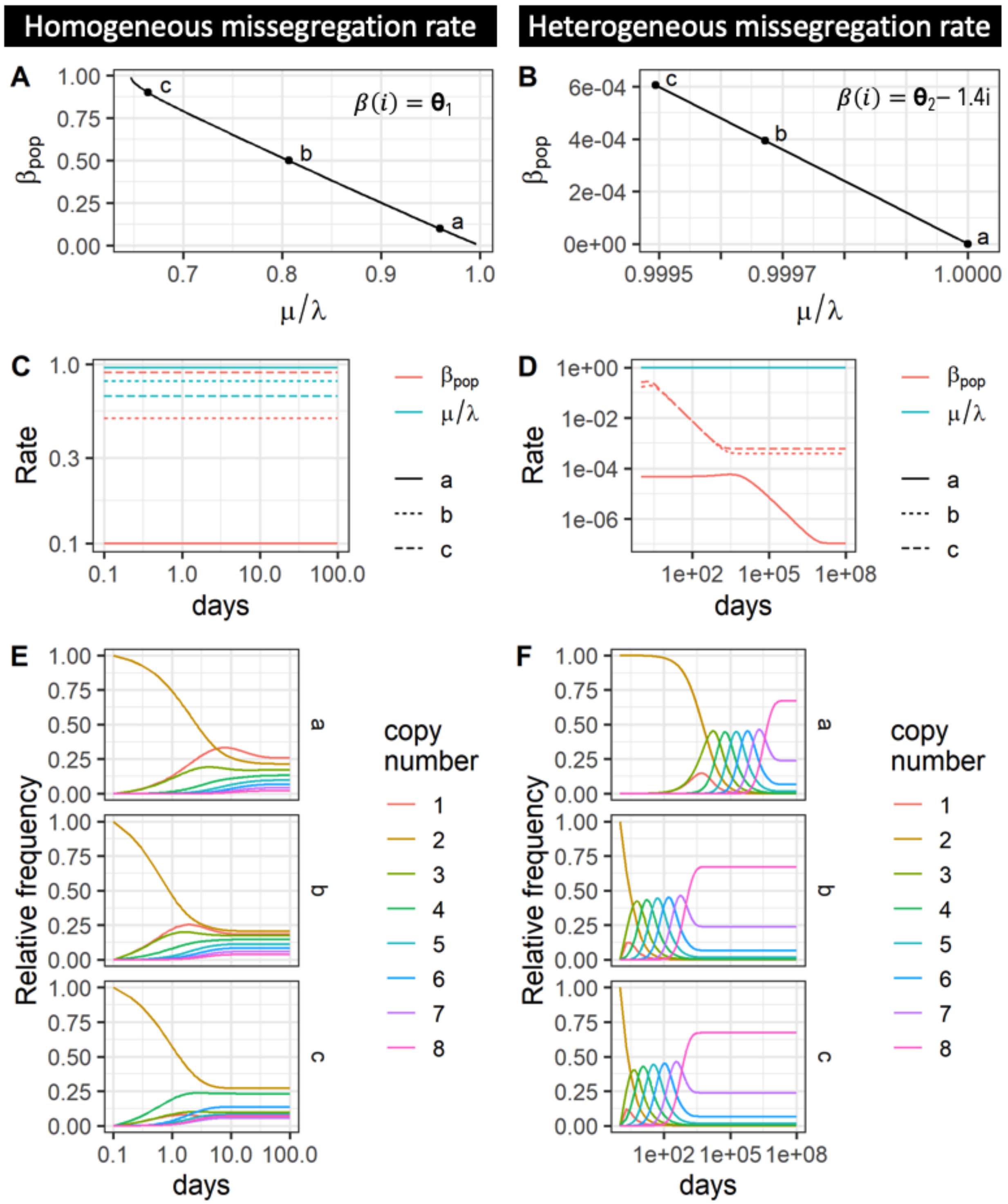


Supplementary Figure 1: **Predicted changes in missegregation rate during tumor evolution.** (A-B) Critical curves displayed as a function of the population average missegregation rate (y-axis) and departure from homeostasis (1 - turnover rate; x-axis). $\lambda := 1$ in all calculations, i.e. cells divide once per day. Copy number ranges from $[1, 8]$ and death rate is independent of copy number ($\mu(i) := M$), whereas missegregation is either also independent (A, $\beta(i) := \Theta_1$) or dependent on copy number (B, $\beta(i) := \Theta_2 - 1.4i$). (C-D) Population average missegregation and turnover rates over time are shown for three parameter combinations $\theta \in a, b, c$ highlighted in (A,B) respectively. (E-F) Evolution of karyotype composition for each of the three parameter combinations is also shown.

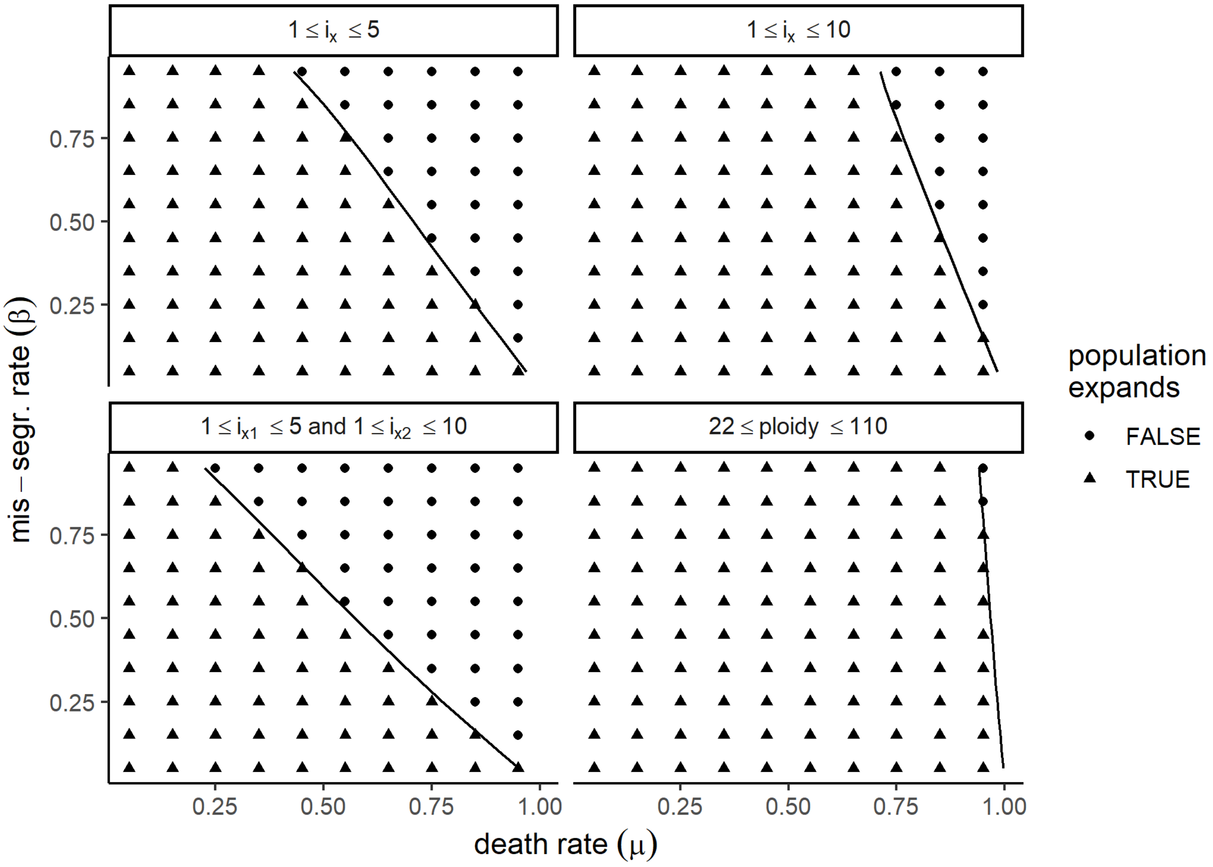


Supplementary Figure 2: **Numerical simulations confirm theoretical MIE curves**. Numerical simulations confirm that the theoretical critical curves shown in Fig. 2A,B separate exponential growth from population extinction.

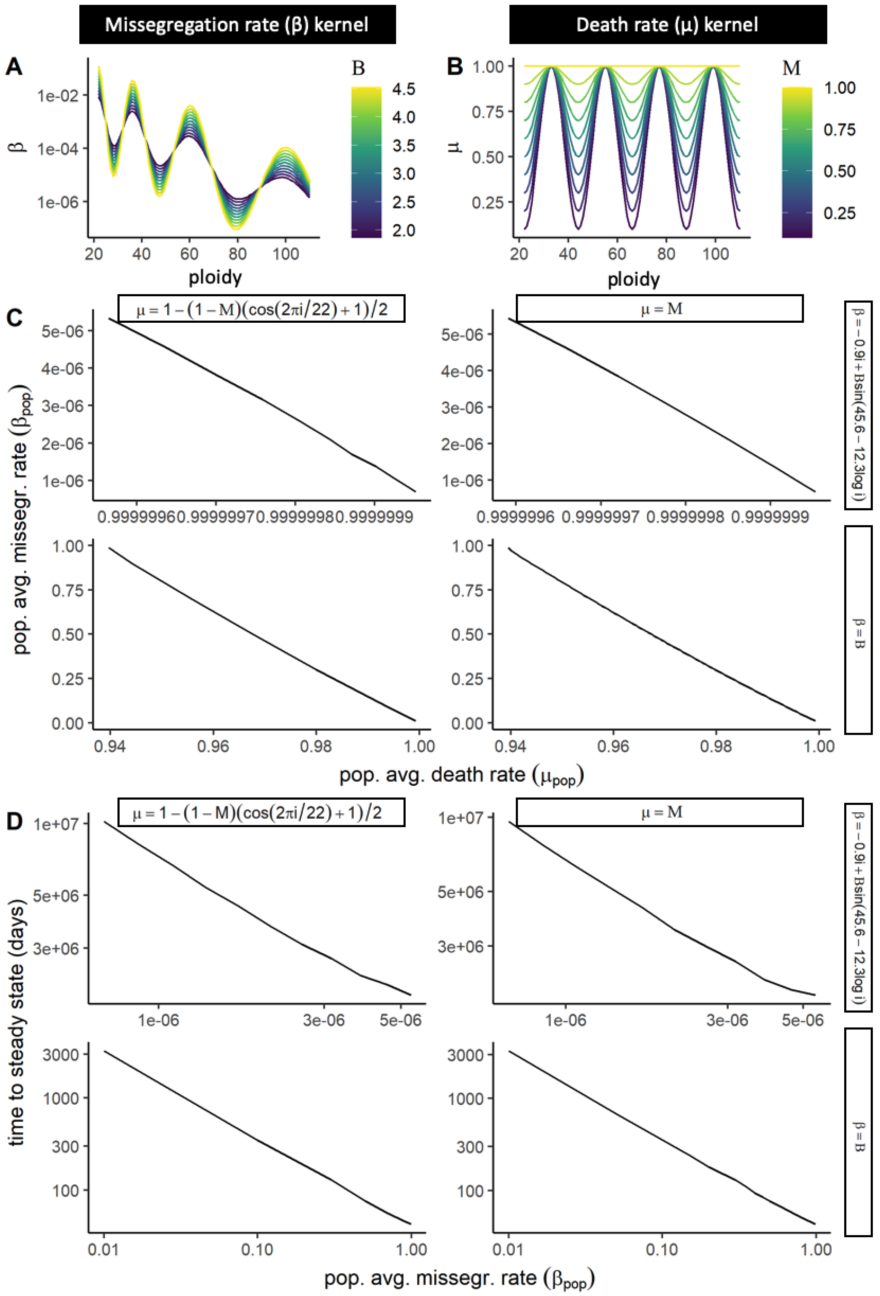


Supplementary Figure 3: **Heterogeneous missegregation and death rates can render MIE impossible**. Kernels used to model intra-tumor heterogeneity in missegregation- (A) and turnover rates (B). (C) Critical curves calculated assuming homogeneous or heterogeneous missegregation and turnover rates are compared with each other. Functions used to model heterogeneous rates are displayed as row and column labels and have ploidy ($i$) as parameter. For each of these functions, we calculated the critical missegregation rate parameter ($B$) and death rate parameter ($M$). Note that for $M = 1$, the turnover rate function in the first column becomes the constant rate from the second column. MIE requires $M \rightarrow 1$, explaining why critical curves look identical in both columns. (D) We also simulated the ODE assuming combinations of the functional forms of missegregation and turnover rates shown in C. Shown here is the number of days needed to reach steady state karyotype composition as a function of population average missegregation rate.

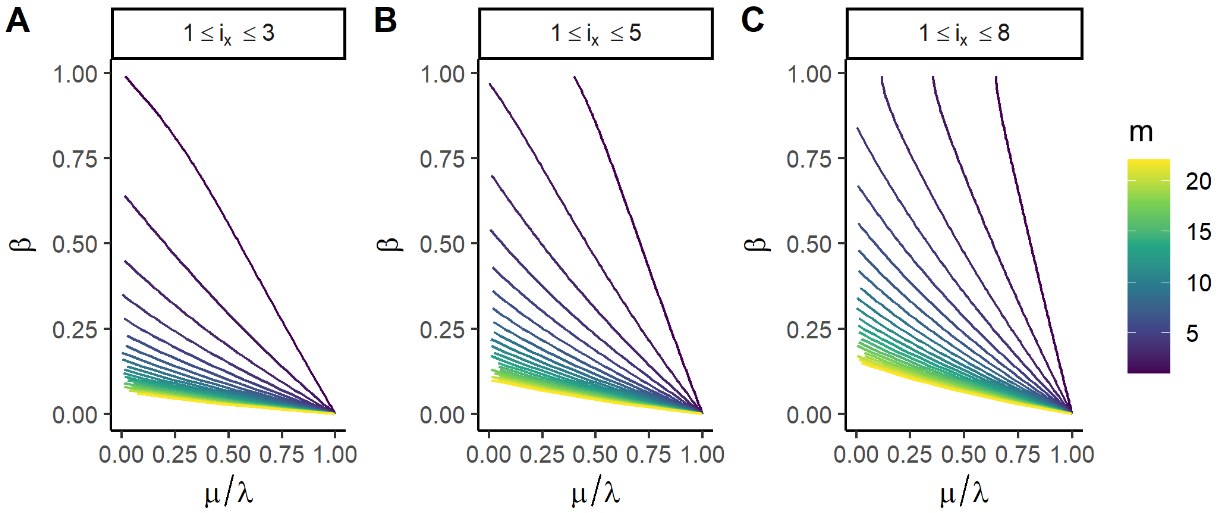


Supplementary Figure 4: **Risk of MIE as a function of the number of chromosome types with finite viable copy number intervals.** (A-C) Critical curves were obtained by finding $(\beta, \frac{\mu}{\lambda})$ for which the maximum eigenvalues of the Jacobian (eq. (9)) is 0. We assume existence of up to $m = 22$ critical chromosomes, where intervals of viable karyotypes for each chromosome type $x \in 1..m$ are defined by $k_x \leq i_x \leq K_x$. We calculate the critical curves assuming cell viability is restricted by all 22 of the chromosomes or by only a subset of them (color code). Hereby we assume each chromosome must have at least one and no more than three (A), five (B) or eight copies respectively (C).

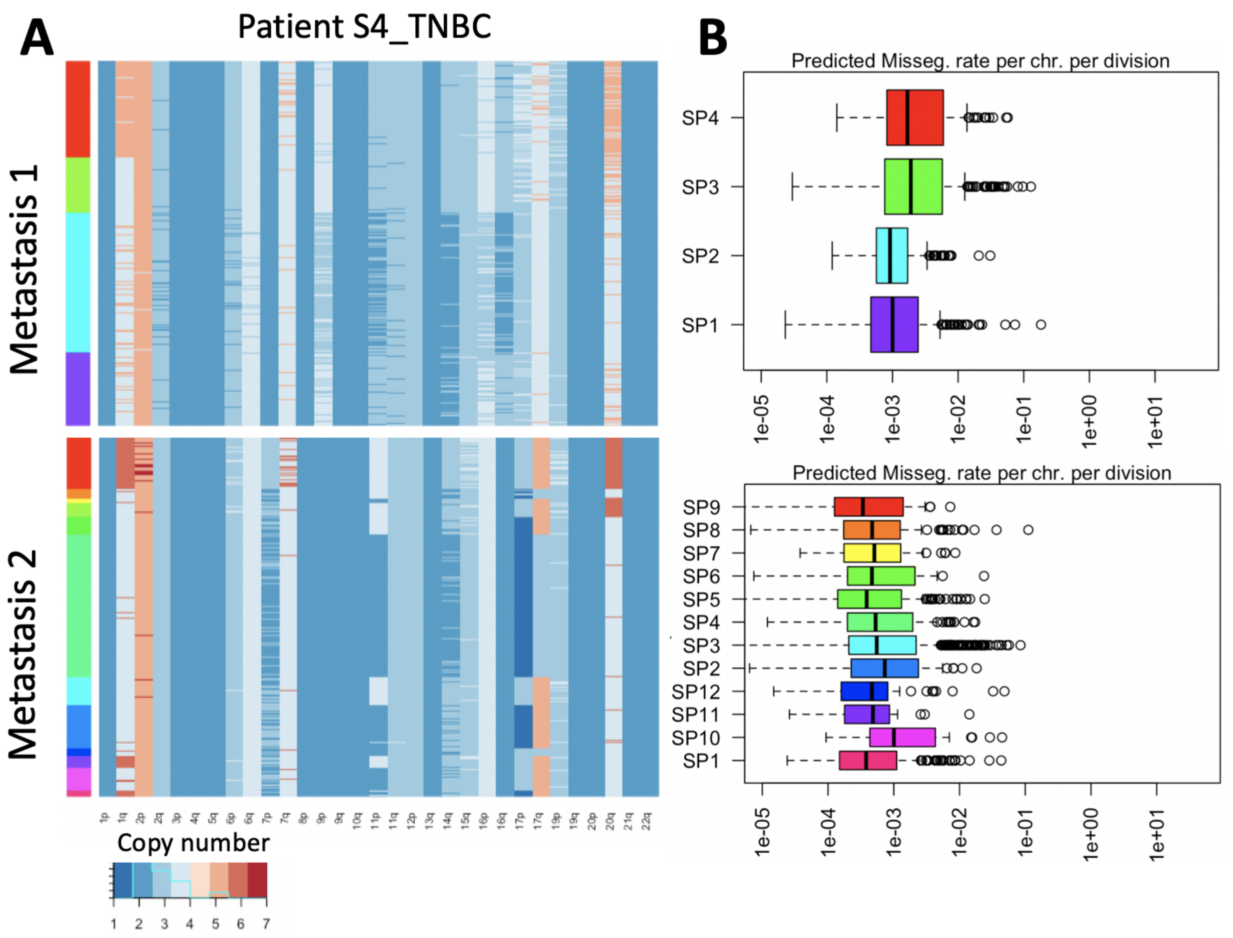


Supplementary Figure 5: **Karyotype profiles inferred for two metastatic TNBC samples .** (A) Copy number profiles from two of the metastatic samples from patient *S4_TNBC1* (marked red in Fig. 3B,D) were clustered into subpopulations of cells with unique karyotypes (left color code). The two metastatic samples originate from the same patient and have significant similarities in their karyotype profiles, even though they were called independently from each other. (B) Inferred mis-segregation rates per cell division per chromosome for each subpopulation in (A).

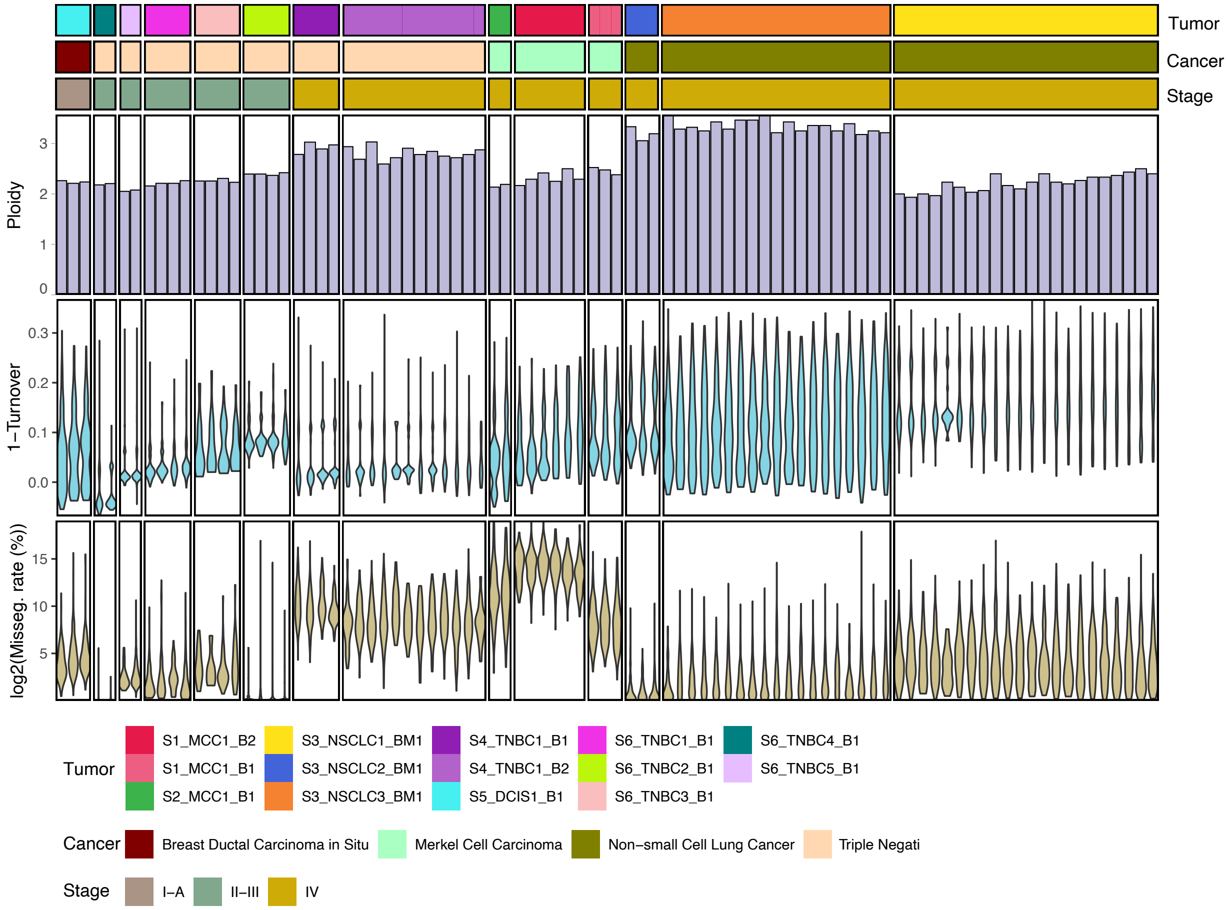


Supplementary Figure 6: **Variability in turnover, missegregation and ploidy across coexisting subpopulations in 14 tumors**. Every column represents a subpopulation of cells with unique karyotype. Subpopulations from the same tumor are grouped together. Their ploidy (A), departure from homeostasis (B) and log2-transformed % missegregation rate (C) are shown.

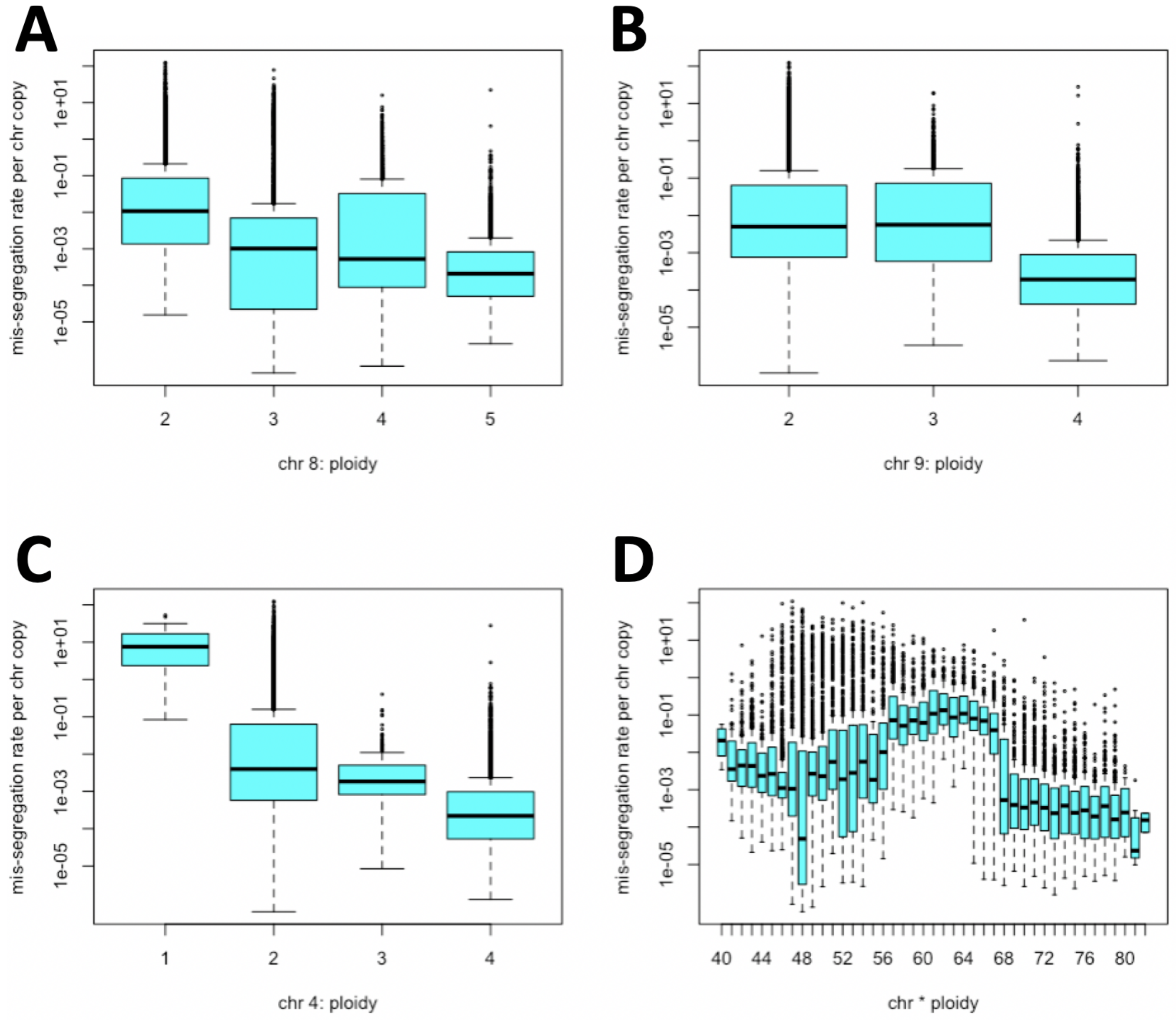


Supplementary Figure 7: **Relationship between karyotype and missegregation rates**. Missegregation rates per chromosome (y-axis) vary with the copy number (x-axis) of specific chromosomes (A-C) and with ploidy (i.e. copy numbers of all chromosomes in aggregate; D).

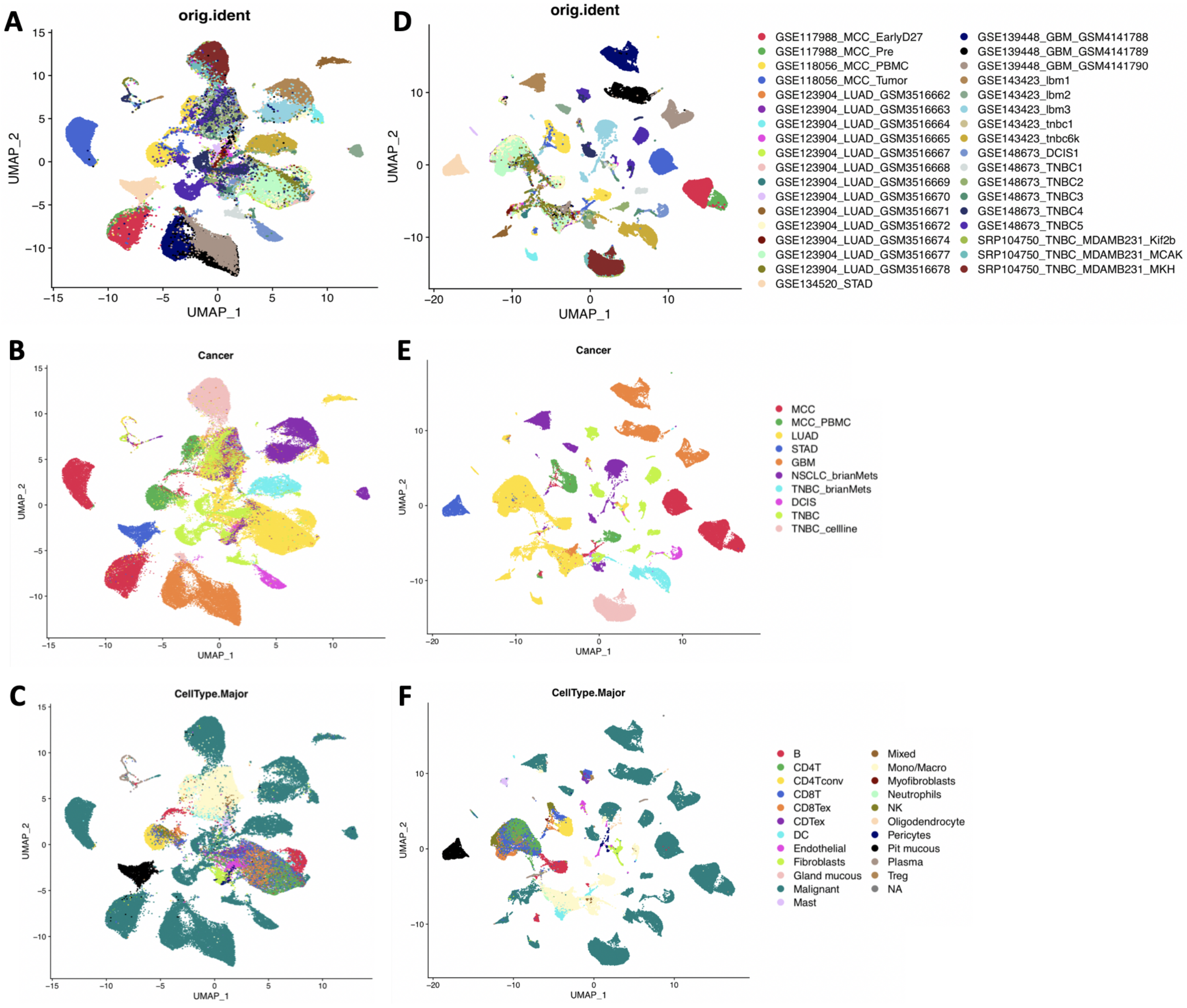


Supplementary Figure 8: **Effect of normalization and scaling for integration of scRNA-seq datasets.** (A-C) UMAP plots using the normalized and scaled data, with the cells labeled by (A) sample name, (B) cancer type of the patients and (C) cell types. Tumor cells cluster by sample origin, and tumor cells from similar cancer types are closer to each other. The PBMC sample of the MCC patient clusters with other normal cells, instead of with the tumor cells from the same patient. Normal cells cluster mostly by cell type. (D-F) Analogous plots to (A-C), but generated using non-normalized data. Generally the cells are more clustered by study, and the tumor cells from different samples are less distinguished, emphasizing the need for normalization and scaling.

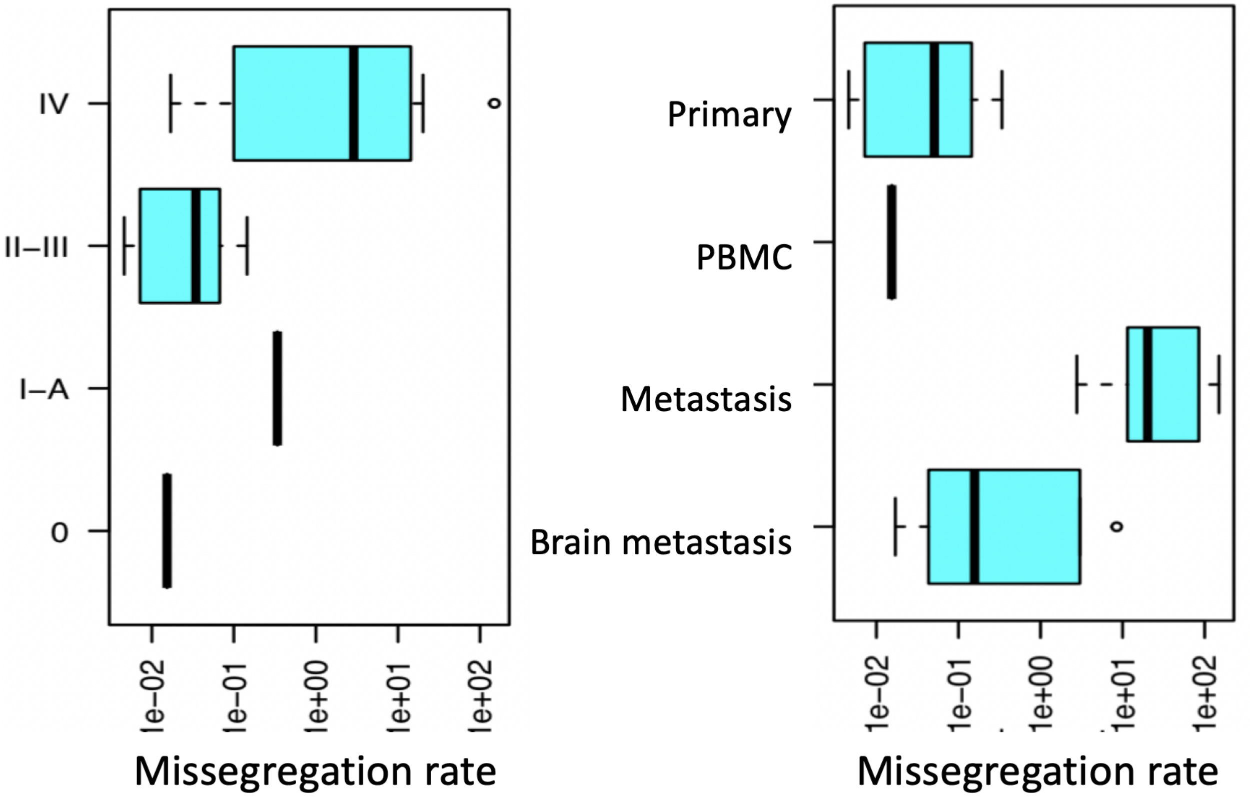


Supplementary Figure 9: **Mis-segregation rates inferred from scRNA-seq derived gene signatures**. Distribution of median mis-segregation rate across all sequenced G1 cells of a given sample are grouped by disease stage (A) or site (B). PBMC:= peripheral blood mononuclear cells.

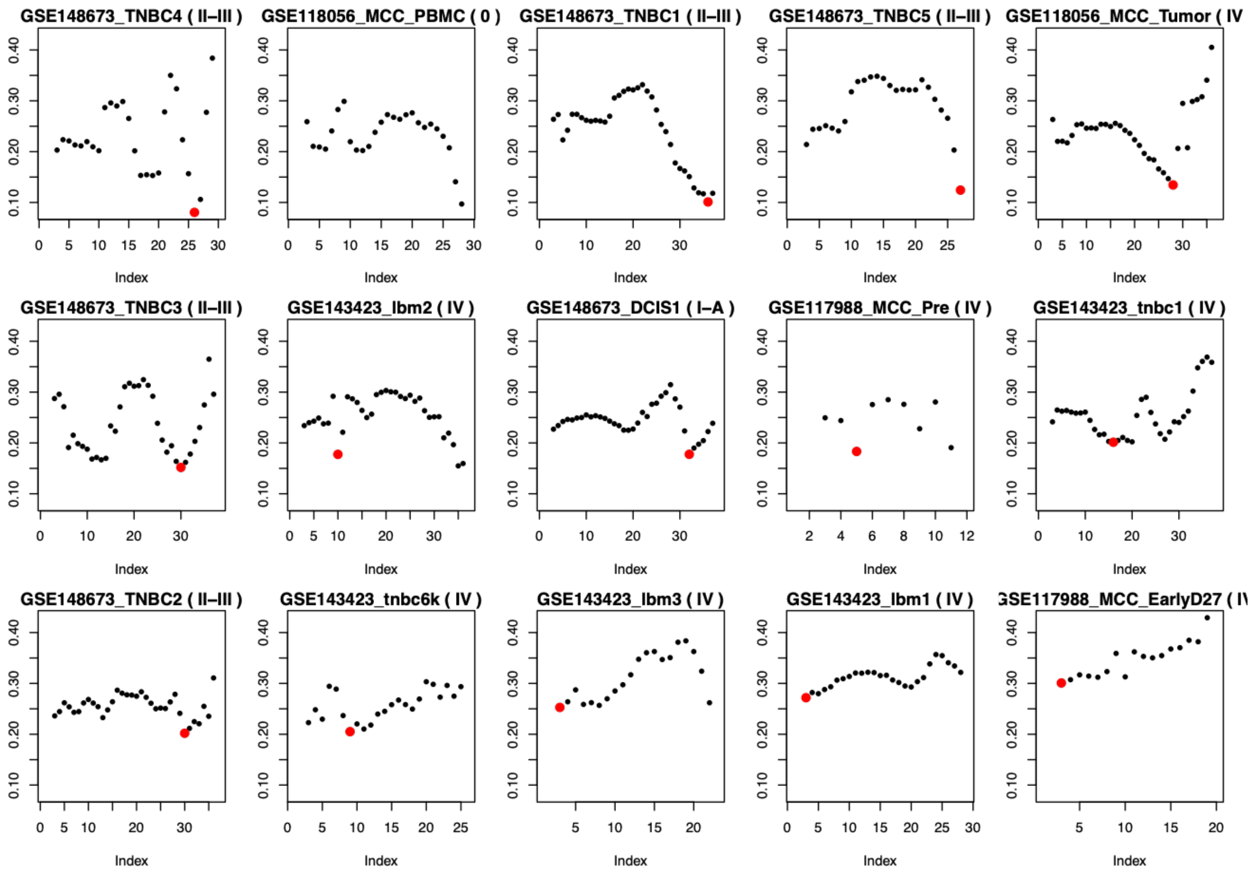


Supplementary Figure 10: **Classification of aneuploid chromosomes with scRNA-Seq derived bias profiles**. Minimum deviation from integer copy numbers (y-axis) guides the choice of x (highlighted red). The more to the right the minimum, the more chromosomes are assumed to be diploid. Shown are the bias profiles for 15 scRNA-seq samples, sorted by global minimum. Samples from late-stage tumors (stage IV) have by trend more non-diploid chromosome arms and their global minimum is higher. Range of x-axis varies because only chromosomes that express a sufficient number of genes in both tumor and normal cells were considered.
